# Noninvasively real-time monitoring in-vivo immune cell and tumor cell interaction by NIR-II nanosensor

**DOI:** 10.1101/2024.12.19.629310

**Authors:** Liwen Huang, Jiang Ming, Zhihua Wang, Jiaxin Wu, Baofeng Yun, Aibin Liang, Yong Fan, Fan Zhang

## Abstract

Immunocytotherapy holds significant promise as a novel cancer treatment, but its effectiveness is often hindered by delayed responses, requiring evaluations every two to three weeks based on current diagnostic methods. Early assessment of immune cell-tumor cell interactions could provide more timely insights into therapeutic efficacy, enabling adjustments to treatment plans. In this study, we report a noninvasive nanosensor (C8R-DSNP) for real-time monitoring of in vivo immune cell activities in the second near-infrared long-wavelength (NIR-II-L) window (1500-1900 nm), which offers deep tissue transparency. The C8R-DSNP responds rapidly to caspase-8, a key apoptotic signaling molecule generated during interactions between natural killer (NK-92) immune cells and tumor cells. Using ratiometric NIR-II-L fluorescence imaging, we captured dynamic in vivo observations of NK-92 cell engagement with tumor cells in a mouse model. These results demonstrated cancer cell apoptosis that happens as early as 4.5 hours after NK-92 cell infusion. Additionally, in vitro urine imaging confirmed the initiation of apoptosis via cleaved fluorescent small molecules, while single-cell tracking within blood vessels and tumors further elucidated immune cell dynamics. This real-time NIR-II-L monitoring approach offers valuable insights for optimizing immunocytotherapy strategies.

## Introduction

Immunocytotherapy is a cancer treatment that involves in vitro activating, engineering, and proliferating immune cells, such as T lymphocytes (T) and natural killer (NK) cells, before infusing them into the patient’s body.^[1]^ A critical aspect of this therapy is the ability of infused NK cells to specifically target and destroy tumor cells through the Fas receptor/Fas ligand (Fas/FasL) pathway^[2]^. In this pathway, FasL on NK cells binds to Fas on cancer cells, triggering the recruitment of Fas-associated death domain (FADD) proteins to initiate “death signals” ^[3]^that result in cancer cell apoptosis and the production of caspase-8.^[4]^ Monitoring caspase-8 activation in real time provides a promising approach to evaluate the immunocytotherapy process. However, due to the delayed response nature of immunocytotherapy, patients often wait weeks to months before assessing treatment efficacy.^[5]^ Real-time monitoring of immune cell-tumor cell interactions could enable earlier evaluation, allowing for timely adjustments to the treatment plan. Since the dynamic interactions between immune and tumor cells require rapid feedback, conventional imaging techniques such as computed tomography (CT), magnetic resonance imaging (MRI), and positron emission tomography (PET) fall short in providing real-time detection with the necessary molecular sensitivity and target specificity.^[6]^ Thus, there is a critical need for advanced visualization tools to accurately and dynamically monitor the immunocytotherapy process, ultimately improving therapeutic outcomes.

To address this challenge, we turned to the optical imaging in the second near-infrared (NIR-II, 1000-1900 nm) window.^[7]^ It has been well-established that NIR-II imaging, particularly in the long-wavelength region (NIR-II-L, 1500-1900 nm),^[8]^ offers superior tissue penetration and higher spatiotemporal resolution, significantly enhancing the ability to detect deep tissue targets noninvasively.^[9]^ Lanthanide-doped downshifting nanoparticles (DSNPs) have garnered considerable attention as promising NIR-II probes due to their unique properties.^[10]^ These nanoparticles, thanks to their multiple energy levels and easily tunable structures, can produce consistent NIR-II-L fluorescence under different excitation wavelengths.^[11]^ This multi-excitation capability enables the development of customized imaging and sensing platforms with real-time feedback.^[12]^ Specifically, using multiple excitation sources for DSNPs allows for more accurate in vivo detection, as one fluorescence signal can serve as a calibration reference, minimizing the impact of signal fluctuations caused by changes in probe concentration during imaging.^[13]^ Activatable DSNPs in the NIR-II-L sub-window have already been utilized for a range of applications, including biological imaging, targeted drug delivery, cancer detection, and treatment monitoring.^[14]^

Here, we report a ratiometric NIR-II-L nanosensor (C8R-DSNP) designed for real-time dynamic monitoring of the immunocytotherapy process, specifically in response to caspase-8 protease, using 808/980 nm excitation. As illustrated in Scheme 1, the C8R-DSNP comprises three essential components: erbium-doped DSNPs, a caspase-8 cleavable substrate, and a renal-cleared ICG molecule (Scheme 1a). Due to the absorption competition-induced emission (ACIE) effect^[15]^, ICG efficiently acts as an excitation filter for the DSNPs at 808 nm resulting in significant quenching of the 1532 nm emission of DSNPs (FL_Ex808 nm_). However, under 980 nm excitation, the 1532 nm emission (FL_Ex980 nm_) remains unaffected. In a mouse tumor model (Scheme 1b), the accumulation of C8R-DSNP at the tumor site can be monitored via FL_Ex980 nm_. Following the infusion of NK-92 cells, caspase-8, an apoptosis initiator in the Fas/FasL signalling pathway, is activated through the interaction of NK-92 cells with cancer cells, cleaving the substrate on the DSNP surface and releasing ICG fragments. This cleavage restores the FL_Ex808 nm_ signal (Scheme 1c). The dynamic process of immunocytotherapy can be assessed in real time by ratiometric NIR-II-L imaging (FL_Ex808 nm_/FL_Ex980 nm_) at the tumor site (Scheme 1d). Additionally, the released ICG fragments are excreted in the urine^[16]^, allowing for urine imaging beyond 850 nm, providing an extra mode for in vitro feedback on the immunocytotherapy process (Scheme 1d). Both in vivo NIR-II-L ratiometric imaging and in vitro urine imaging demonstrate rapid and sensitive detection of caspase-8 activity, with clear signals observed just 4.5 hours after NK-92 infusion. Compared to traditional monitoring methods, which typically take weeks to months, this ratiometric NIR-II-L fluorescence strategy significantly reduces latency, improving efficiency (within 4.5 hours), while avoiding the drawbacks of invasive procedures.

## Results and Discussion

We first synthesized cubic-phase erbium-doped downshifting nanoparticles (DSNP) with a core/multi-shell structure (α-NaYbF_4_:Er,Ce@NaYF_4_:Yb@NaYF_4_:Nd,Yb@NaYF_4_) via a thermal decomposition method^[17]^. The DSNPs could be excited by both 980 nm (absorbed by Yb^3+^) and 808 nm (absorbed by Nd^3+^) excitation light and emit NIR-II-L fluorescence with the peak wavelength of 1532 nm (^4^I_13/2_→^4^I_15/2_) (Figure S1). Ce^3+^ was doped in the core of DSNPs to promote nonradiative relaxation of Er^3+^ (^4^I_11/2_ → ^4^I_13/2_), which enhances the NIR-II-L emission. Considering that the number of sensitizers and Ce^3+^ can have a significant impact on the luminescence, we systematically optimized the doping amounts of Ce^3+^, Yb^3+^ and Nd^3+^ in the core/multi-shell structure (Figure S2-5). The optimized composition for efficient NIR-II-L emission is found to be α-NaYbF_4_:2%Er, 2%Ce@ NaYF_4_:10%Yb@ NaYF_4_:50% Nd, 10%Yb@ NaYF_4_. Then FL_Ex808 nm_ and FL_Ex980 nm_ was further evaluated by adjusting the thickness of the outermost NaYF_4_ shell (Figure S6). The NaYF_4_ shell thickness of 5.5 nm can make the emission intensity of 1532 nm consistent under both 980 nm and 808 nm excitation (Figure S7). The final DSNPs show the core/multi-shell structure and homogeneous spherical shape with the diameter of 44 nm ± 2.3 nm that revealed by the transmission electron microscopy (TEM) images (Figure 1a,b and figure S8). High-resolution TEM images and electron diffractograms of selected regions of DSNPs show the highly crystalline nature (Figure 1b). X-ray diffraction (XRD) measurements confirm the cubic phase of the core/multi-shell structure that consistent with α-NaYF_4_ standard pattern (JCPDS: PDF#39-0724) (Figure S9). We then performed element mapping analysis by scanning the Yb, Nd and Y in the DSNPs, which further demonstrate the core/multi-shell structure with distinct compositional boundaries (Figure S10a). The relative content of elements in different regions of the DSNPs was determined by TEM line scanning. It was evident that Yb and Nd are in the middle layer and Y is distributed in the outermost layer (Figure S10b).

**Figure 1.**
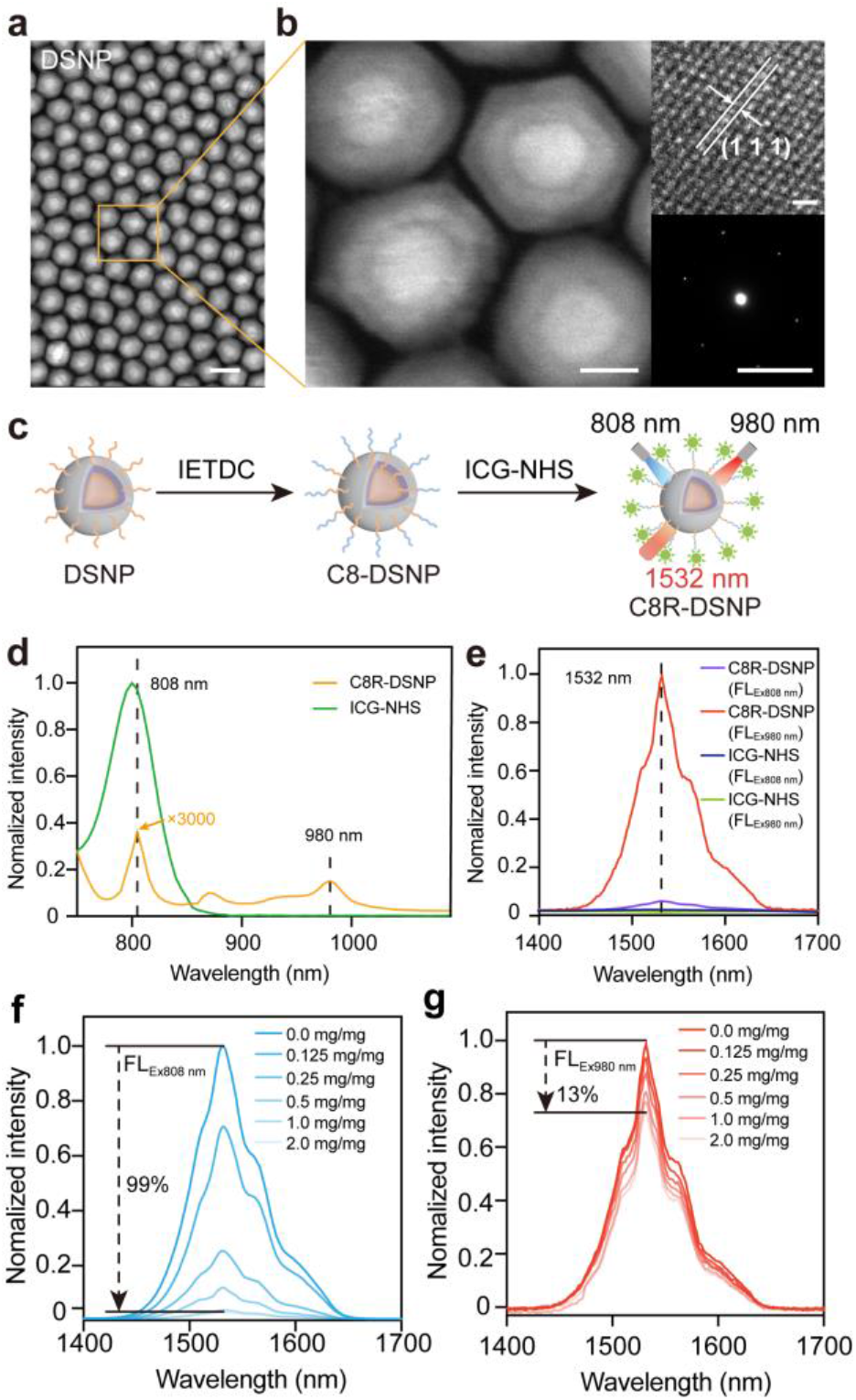
(**a**) Image of dark-field scanning transmission electron microscopy (TEM) of the synthesized DSNPs (scale bar, 50 nm). (**b**) Magnified view of the synthesized DSNP shown in orange rectangle in (**a**) (scale bar, 25 nm). Inset show a high-angle circular dark-field scanning TEM image (top right; scale bar, 10 nm) and the fast fourier transform pattern of the corresponding highresolution image (bottom right; scale bar, 1 nm). (**c**) Schematic illustration of the C8R-DSNP synthesis and ACIE for C8R-DSNP. (**d**) Absorption and emission (**e**) spectra of C8R-DSNP and ICG-NHS. (**f**) FL_Ex808 nm_ and FL_Ex980 nm_ (**g**) intensities of C8R-DSNP after attachment of different concentrations of ICG-NHS.

To synthesize the NIR-II-L nanosensor, peptide substrates IETDC (NH_2_-Ile-Glu-Thr-Asp-Cys-OH) specific for caspase-8 were first synthesized and confirmed by HPLC (Scheme 1a, Figure S11) and LC-MS (Figure S12). The DSNPs were first modified with the amphiphilic polymer COOH-PEG_2000_-Maleimide. Then IETDC was conjugated to DSNPs (C8-DSNP) via sulfhydryl and maleimide reaction. Finally, a modified fluorescent dye ICG (ICG-NHS) was conjugated through EDS-NHS reaction to yield the final nanosensor (C8R-DSNP) (Figure 1c, Figure S13a). After the conjugation of COOH-PEG_2000_-Maleimide, IETDC and ICG, the hydrodynamic diameters of DSNPs increased from 60 ± 2.2 nm, 75 ± 2.4 nm to 94 ± 2.5 nm respectively (Figure S13b,c).The success synthesis of C8R-DSNP can be further verified by Fourier transform infrared spectroscopy (FTIR) (Figure S13d). After the DSNPs were initially conjugated to IETDC, the original carbon-carbon double bond (C=C) in the maleimide breaks, with its characteristic peak at 1650 nm disappearing. Then after modification of ICG, a new characteristic peak of acylamino (C=O) appears at 1700 nm. In addition, the massive zeta potential change from +16.7 ± 1.10 mV (DSNP), −1.21 ± 0.66 mV (C8-DSNP) to −30 ± 2.00 mV(C8R-DSNP) also confirmed the successful surface modification (Figure S13e).

Compared with Nd^3+^ sensitizer which absorbs 808 nm photons, ICG shows a much higher molar extinction coefficient (3000 folds) at this wavelength (Figure 1d). In addition, ICG barely showed absorbance at 980 nm (Figure 1d). Therefore, ICG can serve as the excitation filter for 808 nm and causes a notable fluorescence quenching of C8R-DSNPs at 1532 nm due to ACIE, as compared to that excited under 980 nm excitation (Figure 1e). It should be noted that no emission in the NIR-II-L window was found for ICG under both 808 nm and 980 nm excitation (Figure 1e). To verify the feasibility of the ACIE strategy, we synthesized C8R-DSNPs with different amount (0 to 2 mg/mg) of ICG conjugation. While attenuation of FL_Ex808 nm_ was observed (approximately 99%) with the increase amount of the conjugated ICG (Figure 1f), subtle FL_Ex980 nm_ intensity changes within 13% was determined (Figure 1g). The FL_Ex808 nm_/FL_Ex980 nm_ ratio showed a linear decreased trend with the maximum quenching rate reaching 98% when conjugating with 2.0 mg/mg ICG-NHS (Figure S14). Further increasing the amount of ICG makes no change of FL_Ex808 nm_, which may be due to the saturation of the modification sites on the surface of C8R-DSNP. It should be noted that the excitation filter effect of ICG can only work when ICG molecules were directly conjugated to the DSNPs surface. To verify this, we directly mixed DSNPs with ICG-NHS in trichloromethane solution. As shown in Figure S15, the FL_Ex808 nm_ and FL_Ex980 nm_ only showed 10% decrease with increasing ICG-NHS concentration from 0 to 2 mg/mg.

The response of C8R-DSNPs nanosensor was examined by adding different concentrations of caspase-8 protease. Then NIR-II-L images of the C8R-DSNPs were measured using a commercialized in vivo NIR-II-L imaging system under 808 nm and 980 nm excitation (Figure 2a). As the concentration of caspase-8 increases from 0-1 μM, ICG fragments that released from C8R-DSNPs nanosensor due to caspase-8 cleavage resulted in an apparent recovery of FL_Ex808 nm_ (Figure S16a,b). Since the absorption band of DSNPs at 980 nm is barely affected by the ICG, FL_Ex980 nm_ remained constant (Figure S16c,d). The calculated FL_Ex808 nm_/FL_Ex980 nm_ ratio showed a significant increase with increasing the caspase-8 protease concentration and a linear detectable range in 0-1 μM (Pearson’s coefficient correlation of 0.9183) (Figure 2b,c and Figure S17). We also evaluated the FL_Ex808 nm_/FL_Ex980 nm_ ratio versus the caspase-8 reaction time, the results showed a stably increased ratio with saturation at 30 min (Figure S18), which indicated the fast response of C8R-DSNPs to caspase-8 protease.

**Figure 2.**
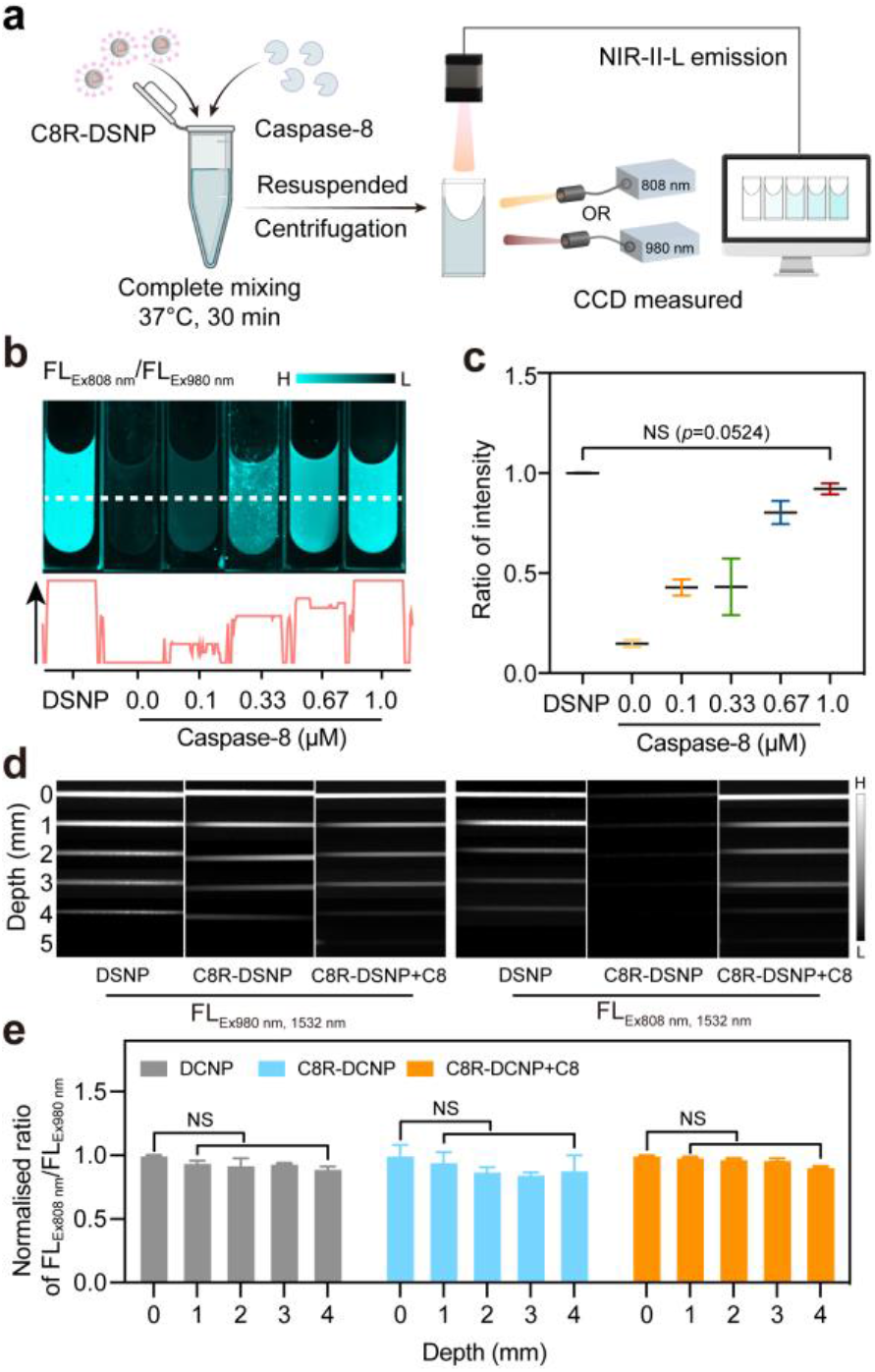
(**a**) Schematic diagram of C8R-DSNPs response to caspase-8 protease based on NIR-II-L imaging. (**b**) Representative NIR-II-L ratiometric fluorescence images and corresponding intensities of C8R-DSNPs over different concentrations of caspase-8 protease. H: high fluorescence intensity; L: low fluorescence intensity. (**c**) Corresponding statistical analysis of normalized FL_Ex808 nm_/FL_Ex980 nm_ ratiometric fluorescence. (**d**) In vitro NIR-II-L imaging of C8R-DSNP filled capillaries before and after caspase-8 response under various mimic tissue depth. (**e**) Statistics on fluorescence ratiometric at different depths in each group of (**d**). H: high fluorescence intensity; L: low fluorescence intensity. Statistical values are expressed as mean ± s.e.m.; NS indicates no significance.

We next adopted a highly scattering medium (1 wt% Intralipid) to mimic the biological tissue for investigating the effect of C8R-DSNPs on caspase-8 protease in deep-tissue. A capillary tube filled with DSNPs, C8R-DSNPs or C8R-DSNPs/caspase-8 mixed solution was immersed in the Intralipid solution at a depth of 0-5 mm. Benefiting from the NIR-II-L emission wavelengths, sharp images were obtained for all the depth (Figure 2d and Figure S19). Although image intensity acquired under 808 nm and 980 nm excitation light for all the three groups decreased with increasing the depth of mimic tissue, the ratiometric fluorescence intensities of FL_Ex808 nm_/FL_Ex980 nm_ showed constant values before and after caspase-8 response (Figure 2e), indicating the reliable results of the ratiometric method in deep tissue and its applicability for in vivo imaging.

The response of the C8R-DSNPs nanosensor to caspase-8 protease activity in living cells was then investigated. First, we evaluated cytotoxicity of C8R-DSNPs by CCK-8 assay using undifferentiated colon cancer cells CT-26, it was found that the C8R-DSNPs were endocytosed by CT-26 cells at 0.5 h post incubation with the maximum phagocytosis at 2 h (Figure S20). No adverse effect was found for the cell viability with C8R-DSNPs concentration up to 500 nM and incubation for 24 h (Figure S21), indicating the well biocompatibility of the C8R-DSNPs nanosensor. After co-culture of CT-26 cells and C8R-DSNPs for 2 h, the IL-2 activated NK-92 cells were added (Figure 3a). It was clearly observed that NK-92 cells (magenta) affixed to the surface of CT-26 cells (green) at 0.5 h post incubation (Figure S22). Then, clusters of NK-92 cells gradually embedded on CT-26 cells and kept unchanged from 1 h-4 h (Figure S22). During this period, NIR-II-L images were taken under both 808 nm and 980 nm irradiation. The increased FL_Ex808 nm_/FL_Ex980 nm_ ratiometric fluorescence also indicate the generation of caspase-8 protease due to interactions of NK-92 cell and CT-26 cell (Figure 3b,c and Figure S23). To further verify that caspase-8 protease indeed generated based on the interaction, enzyme linked immunosorbent assay (ELISA) was carried out to quantitate the interactive product (Figure 3d and Figure S24). The ascending amount of caspase-8 (0-30 ng/mL) versus cell co-culture time correlated well with that from the NIR-II-L ratiometric imaging.

**Figure 3.**
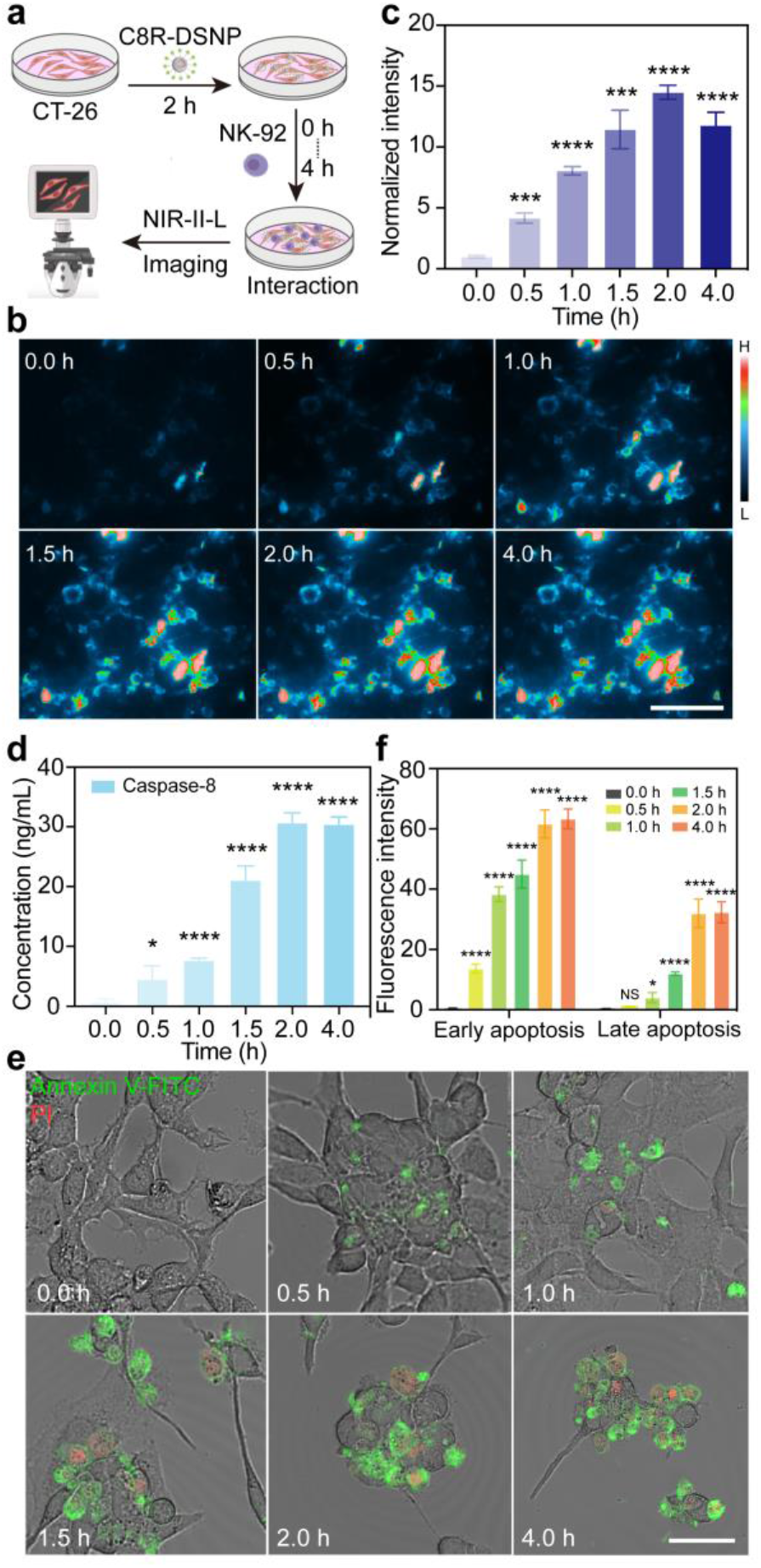
(**a**) Schematic diagram of C8R-DSNP nanosensor responses to caspase-8 protease in live cells. (**b**) Representative NIR-II-L ratiometric fluorescence images of C8R-DSNPs during the interaction of NK-92 with CT-26 cells (scale bar, 10 μm). (**c**) Statistical analysis of NIR-II-L ratiometric fluorescence intensity of C8R-DSNPs at different time points. (**d**) The concentration of caspase-8 protease produced by the interaction of NK-92 and tumor cells versus time. (**e**) Confocal images of cell apoptosis (early apoptosis in green and late apoptosis in red) upon NK-92 cells and CT-26 cells interactions for different time. (**f**) Corresponding fluorescence intensities in (**e**) of early cell apoptosis and late cell apoptosis versus incubation time. Statistically analysand values are shown as mean ± s.e.m.; ^*^P < 0.05, ^***^P < 0.001, ^****^P < 0.0001 (n = 3).

Caspase-8 functions as an initiator of apoptotic signaling and is usually directly associated with early cellular apoptosis^[18]^. To show the therapeutic effect in the cell interaction process, extramembrane phosphatidylserine-binding protein V (Annexin V, early apoptotic cells staining) in combination with propidium iodide (PI, late apoptotic cells staining) were adopted to simultaneously stain the NK-92 treated cancer cells (Figure 3e and Figure S25). The results show that the phenomenon of early cell apoptosis occurred at 0.5 h and continued with treated time (Figure 3f). In the meantime, late apoptotic cells gradually appeared at 1 h due to increased membrane permeability^[19]^. All these results show C8R-DSNPs can still achieve a rapid response to caspase-8 in the relatively complex environment of living cell interactions.

We next adopted CT-26 tumor-bearing mice as the model to demonstrate the ability of C8R-DSNPs for non-invasive monitoring the NK-92 cell therapy (Figure S26a). To determine the optimized time for injection NK-92 and performance of NIR-II-L imaging, we first investigated the accumulation behavior of C8R-DSNPs in tumor of mice (Figure 4a and Figure S26b). We can observe that the tumor vessels were clearly visualized at 0.5 h post injection of the C8R-DSNPs, indicating its fast systematic circulation. Then the tumor margins were gradually lighted up at 2 h and reached maximum intensity at 8 h postinjection (Figure 4b). Due to the enhanced permeation and retention effect (EPR)^[20]^, the tumor intensity then remained stable even for the following 4 h (Figure 4b and Figure S26b), indicating the period when maximum C8R-DSNPs accumulate in the tumor site. The similar imaging procedure was also carried out for NK-92 cells in mice. NK-92 cells pre-labeled with C8R-DSNPs were injected into mice by tail vein, which quickly lighted the whole mice body vessels (Figure 4c and Figure S26c). Driven by the “homing” effect^[21]^, NIR-II-L signals started to show in the spleen and inguinal lymph nodes at 0.5 h post-injection. In the meantime, tumor was also lighted up at 2 h and its intensity gradually increased to the maximum after 4 h through blood circulation (Figure 4d and Figure S26c).

**Figure 4.**
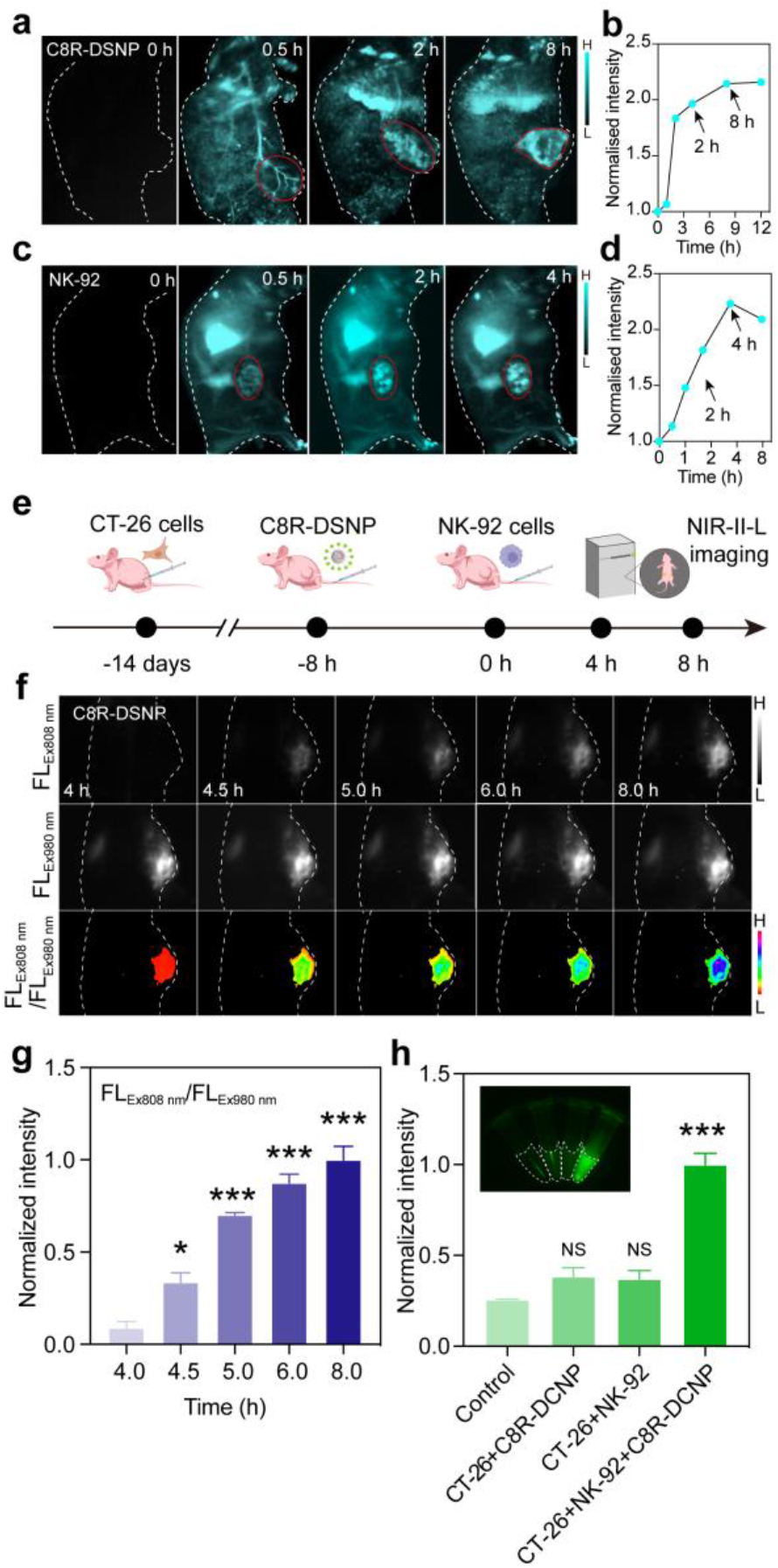
In vivo NIR-II-L imaging of mice at representative time points after injection of C8R-DSNPs (**a**) and NK-92 cells (**c**). Corresponding fluorescence intensity analysis of C8R-DSNPs (**b**) and NK-92 cells (**d**) at the tumor site at different time points. **(e**) Timeline imaging of non-invasive monitoring the NK-92 cell therapy. (**f**) Corresponding NIR-II-L images of mice under 980 nm and 808 nm excitation at different time points (4–8 h). NIR-II-L ratiometric fluorescence results were also shown. H: high fluorescence intensity; L: low fluorescence intensity. (**g**) Statistical analysis of NIR-II-L ratiometric fluorescence intensity of C8R-DSNP at different time points. (**h**) Fluorescence intensities of ICG-fragment in urine from different groups and their statistical analysis. Statistical values are expressed as mean ±s.e.m.; NS indicates no significance, ^*^P < 0.05, ^***^P < 0.001, ^****^P < 0.0001 (n = 3).

Based on the above results, non-invasive monitoring the NK-92 cell therapy was performed by first injection of C8R-DSNPs into the mice and then NK-92 cells 8h later, so that the maximum accumulation of both C8R-DSNP and NK-92 cells could be achieved at the tumor site simultaneously (Figure 4e). Continuous NIR-II-L fluorescence monitoring of tumor sites in mice was obtained at 4 h post injection of NK-92 cells (Video S1). The results showed that the cleaved ICG fragment from C8R-DSNPs makes the NIR-II-L fluorescence recovered at 4.5 h and increased over time due to continuous generation of caspase-8 at the tumor site under 808 nm excitation. (Figure 4f and Figure S27). In the meantime, the production of caspase-8 protease did not affect the NIR-II-L fluorescence under 980 nm excitation (Figure 4f and Figure S27). The FL_Ex808 nm_/FL_Ex980 nm_ ratiometric fluorescence signals were calculated as the feedback of NK-92 cell and tumor cell interaction, which revealed an enhanced therapeutic efficacy with the best performance achieved at 6-8 h for a single time of NK-92 cells transfusion (Figure 4g).

Benefiting from the rapid clearance through kidney, the ICG-peptide fragment can also be adopted as an extra in vitro evaluation way of NK-92 cell and tumor cell interaction (Figure S28). The urine of mice was collected 16 h after injection of NK-92 cells and the strong fluorescence intensity beyond 850 nm can be observed under 760 nm excitation (unable to excite DSNPs) (Figure 4h). In comparison, no florescence signal was observed in healthy mice injected with saline, tumor-bearing mice injected with only C8R-DSNPs or NK-92 cells, which further confirm the successful interaction of C8R-DSNP and NK-92 cells and generation of caspase-8 (Figure 4h).

To further understand the fate of NK-92 cells during immunocytotherapy process in detail, we injected C8R-DSNP-labeled NK-92 cells in to mice bearing CT-26 tumor. Meanwhile, ICG was also injected for highlighting the tumor vascular network under 760 nm excitation to create a reference map around the NK-9 cells (Figure 5a). We monitored their trajectories in real time at the tumor site by recording NIR-II-L microscopic images from the peripheral vasculature to the neovascularisation in the tumor microenvironment as well as to the tumor parenchyma (i-iv in black rectangle in Figure 5b,c). The enrichment of NK-92 cells was firstly observed at the junction of the tumor and the existing peripheral vasculature at 0.5 h post injection (region i). Then they migrated from the tumor margin to the tumor parenchyma through newly formed blood vessels in the next 2-4 h (region ii and iii). With the increase of time, promoted NK-92 cells were traced to the neovascularization intertwined in the tumor site that far from the peripheral vasculature (region iv) (Figure 5c,d), which is consistent with the real-time monitoring of the C8R-DSNPs in wide-field images (Figure 4). Besides, we observed the single NK-92 cell that kept extravasating from the tumor vasculature to reach the tumor parenchyma during the migration process and gradually completing the infiltration process into the tumor tissue (Figure 5c, Figure S29, Figure 5e Video S2). These results will provide clearer insights into the detailed migration, localization and interactions of NK-92 cells at the tumor site and help in understanding how immune cells respond to the tumor microenvironment or other immune challenges.

**Figure 5.**
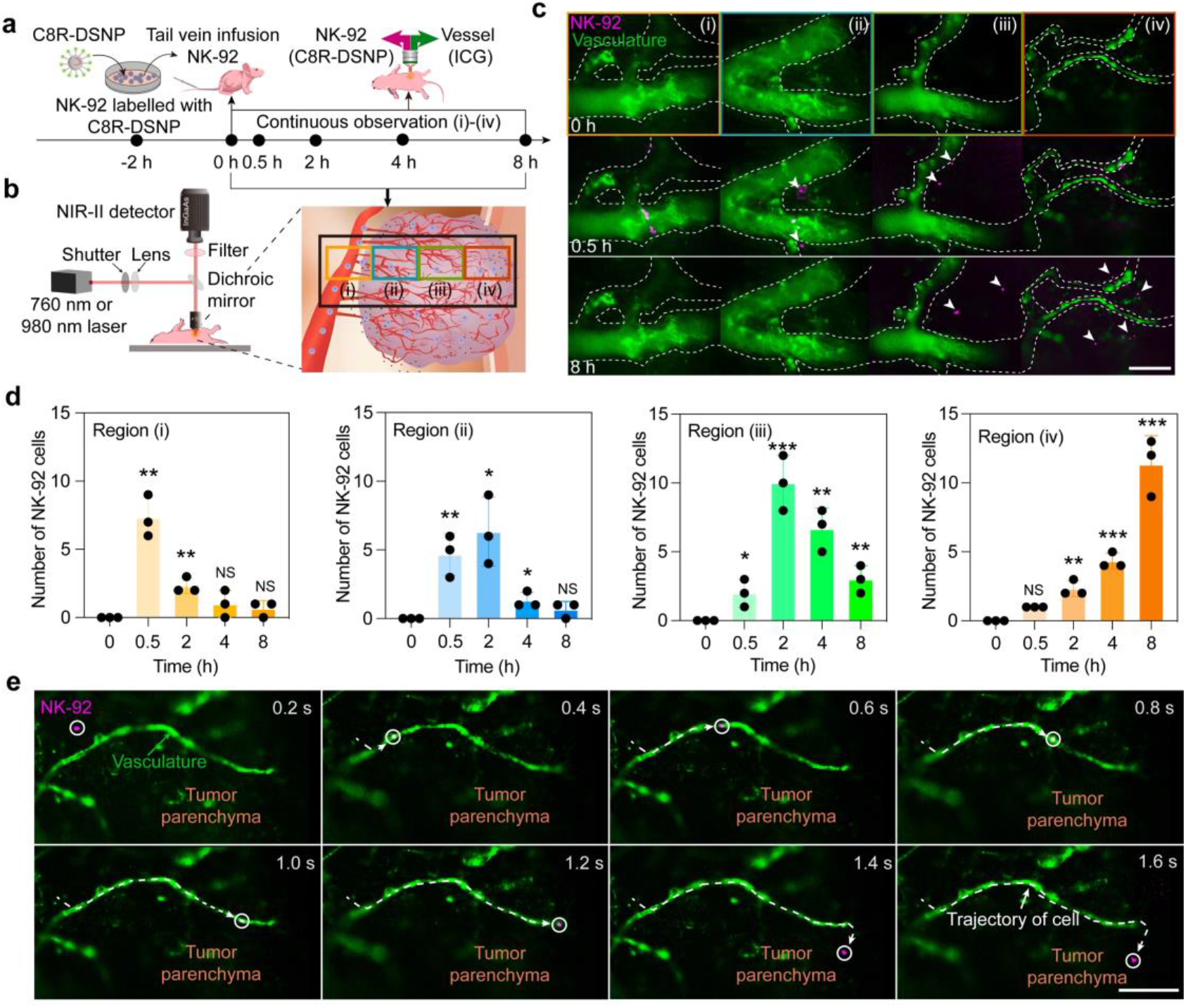
(**a**) Timeline imaging of NK-92 cells in the vasculature of tumour tissue. Blood vessels were visualized by injection of ICG. (**b**) Schematic of in vivo NIR-II-L microscopic imaging of NK-92 cell trajectories in real time at the tumor site. NK-92 cells labelled with DSNPs were injected via tail vein into mice. (**c**) Corresponding multiplexed NIR-II-L images of NK-92 cells (magenta) in the tumor vasculature (green) at different regions and time points. The white arrows point to the extravasated NK-92 cells. Scale bars, 50 μm. (**d**) Statistical analysis of the number of NK-92 cells collected in regions (i)-(iv) versus different time. (**e**) Single NK-92 cell (magenta) trajectory tracking (white dotted line) from the tumor vasculature (green) to the parenchyma. Scale bar, 50 μm. Statistical values are expressed as mean ± s.e.m.; NS indicates no significance, ^*^P < 0.05, ^**^P < 0.01, ^***^P < 0.001 (n = 3).

To further assess the immunocytotherapy of NK-92 cells, we dissected the tumor-bearing mice at different time points. Tumor tissues were extracted and stained with TdT-mediated dUTP nick end labeling (TUNEL) (Figure S30a). The results showed that efficient immunocytotherapy started from 4.5 h after NK-92 injection, which is consistent with the above in vivo imaging results. With the development of immunocytotherapy process, the amount of apoptotic tumor cells (green) and exfoliated tumor parenchyma (white arrows) continued to increase. In the meantime, the proliferation of tumor cells was analyzed, which showed a significantly suppressed trend (Figure S30b).

To evaluate the long-term in vivo toxicity, immunohistochemical assessment and routine blood examination index were performed after tail vein injections of C8R-DSNP nanosensor. The results showed no tissue damage or any other signal of inflammatory lesion as compared to healthy mice (Figure S31). In addition, the number of blood cells and morphological distribution also did not show abnormal indicators (Table S1). This provides a biosafety guarantee for further clinical applications.

## Conclusion

In this study, we reported a ratiometric NIR-II-L nanosensor (C8R-DSNP), which responds to caspase-8 protease, a marker of cell apoptosis. The high signal-to-noise ratio of NIR-II-L fluorescence enables real-time in vivo monitoring of NK-92 cell-cancer cell interactions during immunotherapy, avoiding potential tissue inflammation associated with conventional imaging methods. The C8R-DSNP nanosensor’s real-time feedback capability significantly shortens the evaluation period of NK-92 cell therapy to just 4.5 hours, compared to the weeks or months required by conventional clinical immunotherapy detection cycles. This imaging strategy provides new insights and enhances clinical evaluation of immunotherapy. By conjugating specific intermediate substrates, such as biotin-labeled peptides or small molecule inhibitors^[22]^, the C8R-DSNP platform can be adapted to monitor other disease treatments, such as real-time detection of inflammatory markers in autoimmune therapies^[23]^ or dynamic insulin changes in diabetes management.^[24]^ This versatility not only improves diagnostic accuracy but also opens new possibilities for personalized medicine. Furthermore, incorporating a dual-channel NIR-II-L imaging system^[25]^, in conjunction with quantitative devices like flow analyzers^[26]^, would enable parallel quantitative analysis of immune cells and real-time multiplexed imaging using additional NIR-II-L probes (e.g., Tm^3+^-based DSNPs)^[27]^. This approach would provide a more comprehensive and precise understanding of cell interactions and dynamics during immunotherapy, as well as facilitate more accurate therapeutic evaluations in clinical settings.

## Acknowledgements

Our work was supported from the National Key R&D Program of China (grant no. 2023YFB3507100), and the National Natural Science Foundation of China (NSFC, grant no. 22088101, 22222403, 22161160320, 32171377), and the Research Program of Science and Technology Commission of Shanghai Municipality (grant no. 21142201000, 22JC1400400), and the Innovation Program of Shanghai Municipal Education Commission (2023ZKZD08), and the New Cornerstone Science Foundation through the XPLORER PRIZE.

## Data Availability Statement

The data that support the findings of this study are available in the supporting information of this article.

---

**Scheme 1.**
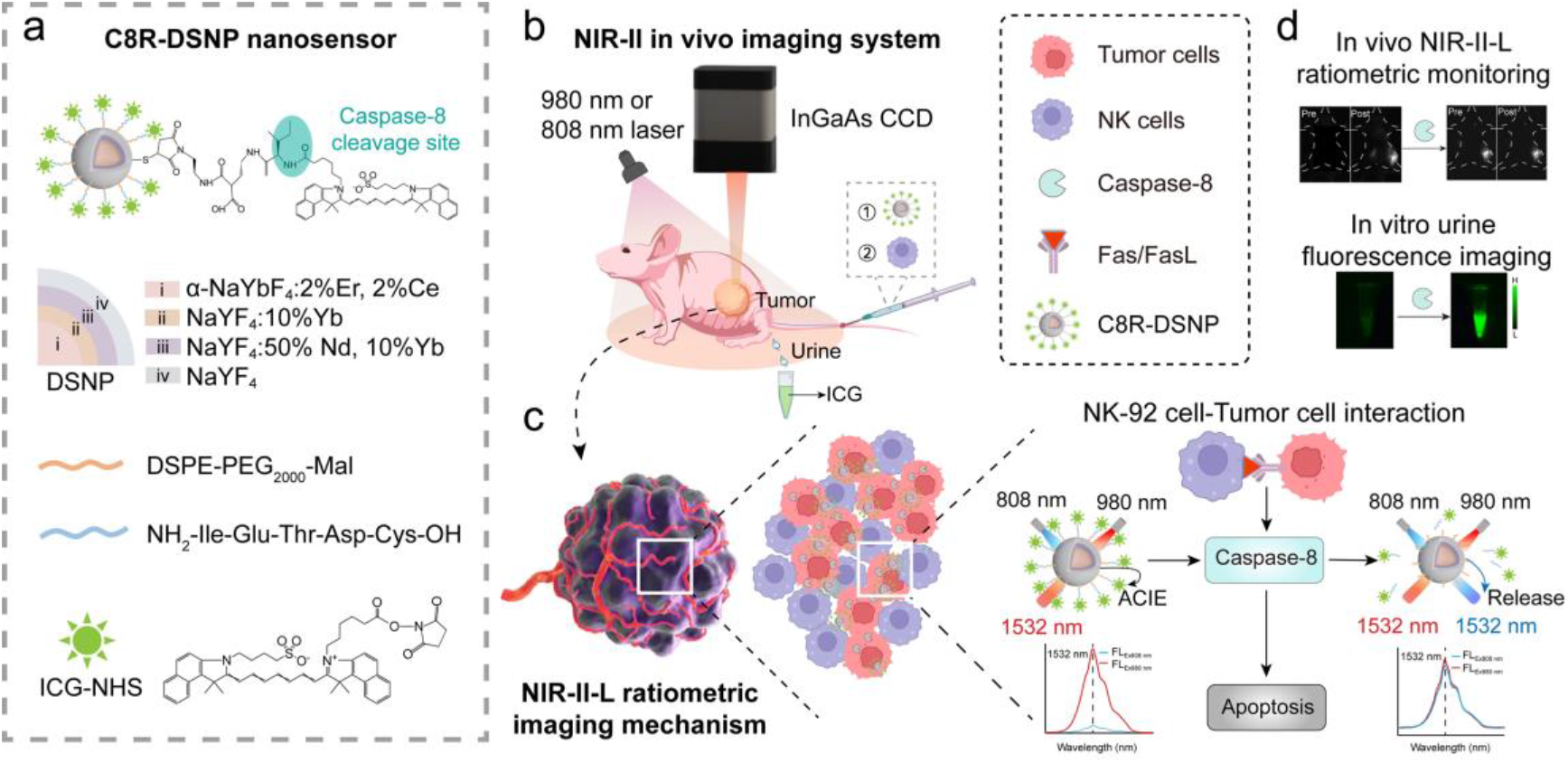
Schematic illustration of non-invasive dynamic monitoring of immunocytotherapy. **a**) DSNP construction and the structure of the C8R-DSNP nanosensor. **b**) Schematic of in vivo non-invasive NIR-II imaging and (**c**) mechanism of NIR-II-L ratiometric fluorescence imaging for NK-92 cell-tumor cell interaction in mice. **d**) Application of non-invasive dynamic monitoring of immunocytotherapy via in vivo ratiometric NIR-II-L imaging and in vitro urinalysis.

