## Supporting Information for "Noninvasively real-time monitoring in-vivo immune cell and tumor cell interaction by NIR-II nanosensor"

Synthesis of the α-$NaYbF_4$:2%Er, 2%Ce@ $NaYF_4$:10%Yb@ $NaYF_4$:50% Nd, 10%Yb nanoparticle: 5

Figure S26 ... 38
Figure S27 ... 39
Figure S28 ... 40
Figure S29 ... 41
Figure S30 ... 42
Figure S31 ... 43
Table S1. ... 44
**References ... 44**
**Author Contributions ... 44**

### Experimental Procedures

#### Chemicals

Peptides were obtained from Synpeptide Co., Ltd (Nanjing, China). caspase-8 protease (mouse) was purchased from MedChemExpress (MCE). Ficoll, Erbium (III) chloride hexahydrate, Yttrium (III) chloride hexahydrate, sodium trifluoroacetate (Na-TFA), 1-octadecene (ODE) and oleic acid (OA) were purchased from Sigma-Aldrich (St Louis, MO, USA). 1-ethyl-3-(3-dimethyl aminopropyl) carbodiimide hydrochloride (EDC • HCl) was purchased from J&K Scientific Ltd. and N-hydroxysuccinimide (NHS) were from Adamas-beta. COOH-PEG2000-Maleimide was purchased from Ponsure biological Co., Ltd (Shanghai, China). Dulbecco’s modified eagle medium (DMEM) and Penicillin–streptomycin (PS) was purchased from Gibco. Hank's balanced salt solution (HBSS), Fetal bovine serum (FBS) was purchased from MesGen Biotech Co., Ltd (Shanghai, China). All chemicals were used as received without any further purification.

#### Characterizations

TEM, high-resolution TEM and high-angle circular dark-field scanning TEM measurements were performed on a JEM-2100F transmission electron microscope equipped with a post-column Gatan imaging filter (GIF-Tridium) at an accelerating voltage of 200 kV. X-ray diffraction measurements were carried out using Cu Kα radiation (wavelength = 1.5406 Å) on a Bruker D8 diffractometer at room temperature. NIR-II-L emission spectra were measured on an Edinburgh FLS980 spectrometer equipped with 980 nm and 808 nm diode lasers (MLL-III-808-2W, Changchun New Industrial Photonics). Absorption spectra were collected by on a PerkinElmer Lambda 750 S UV-Vis-NIR spectrometer. The hydrodynamic radii were obtained by dynamic light scattering measurements (Zetasizer Nano, Malvern Instruments). The NIR-II-L in vivo fluorescence imaging was performed with a modified home-built InGaAs array detector (NIRvana 640, 640 × 512 pixels; Princeton Instruments).

#### Synthesis of DSNP (α-$NaYbF_4$:2%Er, 2%Ce@ $NaYF_4$:10%Yb@ $NaYF_4$:50% Nd, 10%Yb@ $NaYF_4$) core-shell structure

**Synthesis of the α-$NaYbF_4$:2%Er, 2%Ce nanoparticle:** The synthesis of α-$NaYbF_4$:2%Er, 2%Ce nanoparticle is basically the same as that reported in the literature with some modifications. In a typical procedure for the synthesis of α-$NaYbF_4$:2%Er, 2%Ce nanoparticles, 1 mmol of Na-TFA, 0.96 mmol of Yb-TFA, 0.02 mmol of Er-TFA and 0.02 mmol of Ce-TFA were added to a 50 mL flask that contained 10 mL of OA, 10 mL of ODE and 10 mL of OM. The mixture was heated to 140 °C under vacuum until a pellucid solution appeared, the solution was heated to 310 °C at a rate of 15 °C $min^{-1}$, maintained for 20 min under an argon atmosphere and then cooled to room temperature. Nanoparticles were obtained by centrifugation, washed with ethanol three times and redispersed in 10 mL of cyclohexane. The size of nanoparticles is about 7 nm. In a typical procedure for the synthesis of larger α-$NaYbF_4$:2%Er, 2%Ce nanoparticles, 2 mmol of Na-TFA 1.92 mmol of Yb-TFA, 0.04 mmol of Er-TFA, 0.04 mmol of Ce-TFA and 1 mmol α-NaYbF4:2%Er, 2%Ce (7 nm) nanoparticles were added to a 100 mL flask that contained 10 mL of OA and 10 mL of ODE. The mixture was heated to 140 °C under vacuum until a pellucid solution appeared, the solution was heated to 290°C at a rate of 10°C min$^{-1}$, maintained for 40 min under an argon atmosphere and then cooled to room temperature. Nanoparticles were obtained by centrifugation, washed with ethanol three times and redispersed in 10 mL of cyclohexane. The size of nanoparticles is about 12 nm. The synthesis method of α-$NaYbF_4$:2%Er, x%Ce (x = 0, 1, 5, 8 and 10) are the same as that of α-$NaYbF_4$:2%Er, 2%Ce except that Ln-TFA is replaced by the corresponding metal trifluoroacetate. In a typical procedure for the synthesis of largest α-$NaYbF_4$:2%Er, 2%Ce nanoparticles, 4 mmol of Na-TFA 3.84 mmol of Yb-TFA, 0.08 mmol of Er-TFA, 0.08 mmol of Ce-TFA and 1 mmol α-NaYbF4:2%Er, 2%Ce (12 nm) nanoparticles were added to a 100 mL flask that contained 20 mL of OA and 20 mL of ODE. The mixture was heated to 140 °C under vacuum until a pellucid solution appeared, the solution was heated to 290°C at a rate of 10°C min$^{-1}$, maintained for 50 min under an argon atmosphere and then cooled to room temperature. Nanoparticles were obtained by centrifugation, washed with ethanol three times and redispersed in 20 mL of cyclohexane. The size of nanoparticles is about 19 nm. The synthesis method of α-$NaYbF_4$:2%Er, x%Ce (x = 0, 1, 5, 8 and 10) are the same as that of α-$NaYbF_4$:2%Er, 2%Ce except that Ln-TFA is replaced by the corresponding metal trifluoroacetate.

**Synthesis of the α-$NaYbF_4$:2%Er, 2%Ce@ $NaYF_4$:10%Yb nanoparticle:** The synthesis of α-$NaYbF_4$:2%Er, 2%Ce@ NaYF4:10%Yb nanoparticle is basically the same as that reported in the literature with some modifications. In a typical procedure for the synthesis of α-$NaYbF_4$:2%Er, 2%Ce@ $NaYF_4$:10%Yb nanoparticles, 0.2 mmol α-$NaYbF_4$:2%Er, 2%Ce (19 nm), 1.6 mmol of Na-TFA, 0.16 mmol of Yb-TFA and 1.44 mmol of Y-TFA were added to a 100 mL flask that contained 20 mL of OA and 20 mL of OM. The mixture was heated to 140 °C under vacuum until a pellucid solution appeared, the solution was heated to 290 °C at a rate of 10 °C min$^{-1}$, maintained for 50 min under an argon atmosphere and then cooled to room temperature. Nanoparticles were obtained by centrifugation, washed with ethanol three times and redispersed in 10 mL of cyclohexane. The synthesis method of α-$NaYbF_4$:2%Er, x%Ce@ $NaYF_4$:10%Yb (x = 0, 1, 5, 8 and 10) and α-$NaYbF_4$:2%Er, 2%Ce@ $NaYF_4$:y%Yb (y = 0, 5, 30, 50 and 70) are the same as that of α-$NaYbF_4$:2%Er, 2%Ce@ $NaYF_4$:10%Yb except that Ln-TFA is replaced by the corresponding metal trifluoroacetate.

**Synthesis of the α-$NaYbF_4$:2%Er, 2%Ce@ $NaYF_4$:10%Yb@ $NaYF_4$:50% Nd, 10%Yb nanoparticle:** The synthesis of α-$NaYbF_4$:2%Er, 2%Ce@ $NaYF_4$:10%Yb@ $NaYF_4$:50% Nd, 10%Yb nanoparticle is basically the same as that reported in the literature with some modifications. In a typical procedure for the synthesis of α-$NaYbF_4$:2%Er, 2%Ce@ $NaYF_4$:10%Yb@ $NaYF_4$:50% Nd, 10%Yb nanoparticles, 0.1 mmol α-$NaYbF_4$:2%Er, 2%Ce@ $NaYF_4$:10%Yb, 1.6 mmol of Na-TFA, 0.16 mmol of Yb-TFA, 0.16 mmol of Yb-TFA and 0.64 mmol of Y-TFA were added to a 100 mL flask that contained 20 mL of OA and 20 mL of OM. The mixture was heated to 140 °C under vacuum until a pellucid solution appeared, the solution was heated to 280 °C at a rate of 10 °C $min^{-1}$, maintained for 70 min under an argon atmosphere and then cooled to room temperature. Nanoparticles were obtained by centrifugation, washed with ethanol three times and redispersed in 10 mL of cyclohexane. The synthesis method of α-$NaYbF_4$:2%Er, x%Ce@ $NaYF_4$:10%Yb (x = 0, 1, 5, 8 and 10)@ $NaYF_4$:50% Nd, 10%Yb, α-$NaYbF_4$:2%Er, 2%Ce@ $NaYF_4$:y%Yb (y = 0, 5, 30, 50 and 70)@ $NaYF_4$:50% Nd, 10%Yb, α-$NaYbF_4$:2%Er, 2%Ce@ $NaYF_4$:10%Yb@ $NaYF_4$:50% Nd, z%Yb (z = 0, 4, 7, 30 and 50) and α-$NaYbF_4$:2%Er, 2%Ce@ $NaYF_4$:10%Yb@ $NaYF_4$:p% Nd, 10%Yb (p = 0, 10, 30, 50 and 90) are the same as that of α-$NaYbF_4$:2%Er, 2%Ce@ $NaYF_4$:10%Yb@ $NaYF_4$:50% Nd, 10%Yb except that Ln-TFA is replaced by the corresponding metal trifluoroacetate.

**Synthesis of the α-$NaYbF_4$:2%Er, 2%Ce@ $NaYF_4$:10%Yb@ $NaYF_4$:50% Nd, 10%Yb@ $NaYF_4$ nanoparticle:** The synthesis of α-$NaYbF_4$:2%Er, 2%Ce@ $NaYF_4$:10%Yb@ $NaYF_4$:50% Nd, 10%Yb@ $NaYF_4$ nanoparticle is basically the same as that reported in the literature with some modifications. In a typical procedure for the synthesis of α-$NaYbF_4$:2%Er, 2%Ce@ $NaYF_4$:10%Yb@ $NaYF_4$:50% Nd, 10%Yb@ $NaYF_4$ nanoparticles, 0.1 mmol α-$NaYbF_4$:2%Er, 2%Ce@ $NaYF_4$:10%Yb@ $NaYF_4$:50% Nd, 10%Yb, 3.2 mmol of Na-TFA and 3.2 mmol of Y-TFA were added to a 100 mL flask that contained 20 mL of OA and 20 mL of OM. The mixture was heated to 140 °C under vacuum until a pellucid solution appeared, the solution was heated to 290 °C at a rate of 10 °C $min^{-1}$, maintained for 70 min under an argon atmosphere and then cooled to room temperature. Nanoparticles were obtained by centrifugation, washed with ethanol three times and redispersed in 10 mL of cyclohexane. The synthesis method of α-$NaYbF_4$:2%Er, x%Ce@ $NaYF_4$:10%Yb (x = 0, 1, 5, 8 and 10)@ $NaYF_4$:50% Nd, 10%Yb@ $NaYF_4$, α-$NaYbF_4$:2%Er, 2%Ce@ $NaYF_4$:y%Yb (y = 0, 5, 30, 50 and 70)@ $NaYF_4$:50% Nd, 10%Yb@ $NaYF_4$, α-$NaYbF_4$:2%Er, 2%Ce@ $NaYF_4$:10%Yb@ $NaYF_4$:50% Nd, z%Yb (z = 0, 4, 7, 30 and 50) @ $NaYF_4$ and α-$NaYbF_4$:2%Er, 2%Ce@ $NaYF_4$:10%Yb@ $NaYF_4$:p% Nd, 10%Yb (p = 0, 10, 30, 50 and 90)@ $NaYF_4$ are the same as that of α-$NaYbF_4$:2%Er, 2%Ce@ $NaYF_4$:10%Yb@ $NaYF_4$:50% Nd, 10%Yb@ $NaYF_4$. The synthesis method of α-$NaYbF_4$:2%Er, 2%Ce@ $NaYF_4$:10%Yb @ $NaYF_4$:50% Nd, 10%Yb@ $NaYF_4$ with different shell thickness are the same as that of α-$NaYbF_4$:2%Er, 2%Ce@ $NaYF_4$:10%Yb@ $NaYF_4$:50% Nd, 10%Yb@ $NaYF_4$ except change the amount of Y-TFA.

#### Surface modification of water-soluble DSNP

Hydrophobic nanoparticles were transferred into the aqueous phase: DSNP nanoparticles (0.1 mM) were transferred into a round-bottom flask and mixed with COOH-PEG2000-Maleimide (25 mg/mL) in a 1:5 ratio by volume, after which trichloromethane was added sequentially to dissolve the particles up to 5 mL. Gradient spinning (-500 kpa, -250 kpa, -150 kpa, -50 kpa, 0 kpa) was performed at room temperature for 1 h. After that, the nanoparticles were placed in a 100 °C oven for 5 min, and then removed and ultrasonicated in a water bath at 100 °C for 30 s. During the process of ultrasonication, 4 mL of ultrapure water at 100 °C was added to dissolve the nanoparticles. Finally, the nanoparticles were centrifuged at high speed (20,000 rpm, 15 min), dispersed in aqueous solution and stored at -20 °C for further use.

#### Synthesis of C8R-DSNP nanosensor

We would like to thank to Dr X.L. form our lab for providing the ICG-COOH. further ICG-COOH will be transformed into ICG-NHS for the next conjugate linkage experiments. Specifically, DSNPs were first mixed with the peptide substrate of caspase-8, NH2-Ile-Glu-Thr-Asp-Cys-OH (IETDC), at a molar ratio of 1:1 in Tris buffer (pH 8.5), and stored overnight at room temperature protected from light. Then tris(2-carboxyethyl)phosphine (TCEP, 1 mM) was added and stirred for 2 h to obtain DSNP-IETDC . The DSNP obtained in the previous step was then initially dissolved with 5 mg of ICG-COOH, 30 mg of 1-ethyl-(3-dimethylaminopropyl) carbodiimide hydrochloride (EDC-HCl), and 30 mg of N-hydroxybutanediimine (NHS) in 5 mL of DMF. The solution was then added to 5 mL of deionised water (pH 6.5) containing 2-(N-morpholino)ethanesulfonic acid (MES) and stirred overnight. After filtration and drying, excess cross-linker was removed and DSNP-IETDC-ICG (C8R-DSNP) was dispersed in an aqueous solution and stored at -20 °C for backup.

The labeling ratio (R) for C8R-DSNP nanosensor was determining by individually counting the amounts of ICG molecules and C8R-DSNP in the purified solution. The number of C8R-DSNP (NP) in solution was counted by Nanoparticle Tracking Analysis (NTA) on a Malvern NanoSight NS300 system, while the molar concentration of ICG ($C_{ICG}$) in the solution was measure by UV absorption spectrum (R1). The equation for calculating labeling ratio was $R=(A_{ICG}/\varepsilon/b*V)/ (NP /NA*V*D)$; Where the $A_{ICG}$ is the absorbance of ICG molecule in solution; $\varepsilon$ is the extinction coefficient of ICG; V is the volume of sample; NP is the counted number of C8R-DSNP; NA is the Avogadro constant; D is the dilution ratio when performing NTA. The labeling ratio was calculated to be around 300 ± 10 ICG molecules per C8R-DSNP.

#### In vitro characterization of caspase-8 protease response by C8R-DSNP nanosensors

C8R-DSNP was incubated with different concentrations of caspase-8 (0 μM, 0,1 μM, 0.33 μM, 0.67 μM, 1.0 μM) in saline at 37 °C for 30 min, or with 0.67 μM caspase-8 in saline at 37 °C for different times (0 min, 10 min, 30 min, 60 min, 120 min), then the mixed solution was centrifuged (20000 rpm, 5 min) and the precipitate was dissolved in decolonized water. The CCD fluorescence intensity of the above samples was characterized using 808 nm and 980 nm lasers, and the filters used were 1300 nm LP and 1400 nm LP.

#### Tissue phantom imaging study

Here 1% Intralipid was chosen as a simulated tissue as it has similar optical properties compared to biological tissues. Glass capillaries were filled with aqueous DSNP, C8R-DSNP and encapsulated C8R-DSNP after caspase-8 cleavage at different concentrations for imaging. The capillaries were then placed under the cylindrical culture dish and covered with different volumes of 1% fat emulsion.

CCD imaging was performed using 808 nm and 980 nm lasers, and the filters used were 1300 nm LP and 1400 nm LP.

#### Cell culture

Natural killer cells (NK-92, Cat No.: CC-Y1400) were provided by Shanghai Yuchi Biochemical Technology Co., Ltd. NK-92 cells were cultured using MEMα complete medium containing 12.5% horse serum (HS), 12.5% fetal bovine serum (FBS), 0.2 mM inositol, 0.1 mM 2-mercaptoethanol, 0.02 mM folic acid and 1% penicillin/streptomycin (PS). NK-92 cell activity was continuously induced by the addition of 200 U/mL recombine IL-2. NK-92 cells were maintained in suspension, with most cells aggregated into clusters and a few dispersed. Colorectal cancer cells (CT-26, Cat No.: TCM37) were provided by Shanghai Bihe Biochemical Technology Co., Ltd. CT-26 cells were cultured using RPMI-1640 complete medium containing 10% FBS and 1% PS. The above cell lines were cultured in an incubator at 37 °C with 5% $CO_2$ and 95% air.

#### Endocytosis assay and cytotoxicity assay of C8R-DSNP

500 μM of C8R-DSNP was added to CT-26 cells (1 × $10^6$) for incubation. The $FL_{Ex980\ nm}$ fluorescence intensity of intracellular C8R-DSNP was observed by NIR microimaging at different time points to determine the optimal time point for C8R-DSNP to be endocytosed by cells. The effect of C8R-DSNP on cytotoxicity was detected by Cell Counting Kit-8 kit (cck-8). CT-26 cells were cultured in 96-well plates for 12 h. After culturing until the cells were well adhered to the wall, different concentrations of C8R-DSNP (0 nM, 10 nM, 50 nM, 100 nM, 200 nM, 500 nM) were added, and the culture was continued for 24 h. The supernatant was discarded and washed with 1 × PBS, and then fresh medium (100 μL/well) was added. After that, 10 μL of cck-8 solution was added to each well and the incubation was continued in the cell culture incubator for 1 hr. Absorbance was measured at 450 nm using an enzyme marker assay. 500 nM C8R-DSNP was added, and the above steps were repeated, and the absorbance was measured at different time points (0 h, 1 h, 2 h, 4 h, 12 h, 24 h).

#### Scanning electron microscopy (SEM) experiment

The samples were fixed with 4% paraformaldehyde and then dehydrated in a gradient with ethanol (30%, 50%, 70%, 90%, 100%). Afterwards, the samples were placed in a freeze-drying oven for overnight freeze-drying. Since biological tissues are not electrically conductive, they need to be gold-spray coated to increase the topographical lining prior to SEM observation and image acquisition was performed by Gemini Zeiss SEM500.

#### In vitro fluorescence imaging of C8R-DSNP nanosensor in the cell-cell interaction.

CT-26 cells (1 × $10^5$) were cultured in confocal culture dish for 24 h. 1 × PBS was washed and C8R-DSNP nanosensor was added after 2 h of incubation. The medium was then discarded, washed with 1 × PBS, and NK-92 cells (1 × $10^5$) were added for co-culture at 37 °C. NIR-II-L fluorescence images were collected using an Olympus-IX71 fluorescence microscope under 808 nm and 980 nm excitation (200 kW $cm^{-2}$, exposure time: 200 ms per frame). Fluorescence images were collected with 1300 LP and 1400 LP.

#### Apoptosis assay

Aspirate the cell culture solution, add 1 × PBS and wash once, add 195 μl Annexin V-FITC conjugate and 5μl Annexin V-FITC, mix gently. Afterwards, add 10μl of propidium iodide staining solution and mix gently. Incubate for 10-20 min at room temperature (20-25 ºC) away from light. CT-26 cells were then subjected to in situ fluorescence detection in a confocal fluorescence microscope, with Annexin V-FITC fluorescing in green and propidium iodide (PI) fluorescing in red.

#### ELISA experiment

NK-92 cells were co-cultured with CT-26 cells for different time points (0 h, 0.5 h, 1 h, 1.5 h, 2 h, and 4 h). After the incubation period, the original medium was discarded, and the cells were washed with PBS. The cells were lysed by repeated freeze-thaw cycles, and the lysate was centrifuged at 1000 rpm for 10 min. The supernatant was collected, and the samples were diluted according to the instructions of the ELISA kit (Bioswamp). Standard samples and experimental samples were added to the wells of a microtiter plate, with each standard and sample replicated three times. The standards were used to generate a standard curve. The plate was incubated at room temperature for 1-2 h. After incubation, the plate was washed multiple times with wash buffer to remove unbound proteins and antibodies. Enzyme conjugates were then added to each well, followed by a second incubation. The plate was washed again to remove excess conjugates. Next, the substrate solution (TMB) was added, and the plate was incubated until a blue color developed. A stop solution, usually dilute sulfuric acid, was then added to stop the reaction, turning the blue color to yellow. The optical density (OD) was measured using a microplate reader at 450 nm. A standard curve was constructed using the data from the standards, with the x-axis representing the concentration of the standard and the y-axis representing the OD values. The caspase-8 concentration in the samples was determined by comparing their OD values to the standard curve.

#### Animal

All animal experimental protocols were conducted in strict accordance with the National Institutes of Health Guide for the Care and Use of Laboratory Animals and approved by the Animal Care and Use Committee of Fudan University. All animal experiments were authorized by the Shanghai Municipal Commission of Science and Technology. Balb/c nude mice (4-6 weeks old, female, average weight ~20 g) were purchased from Shanghai JSJ Laboratory Animal Co. The ambient relative humidity was 55-65% and the temperature was ~25 °C. All mice were anesthetized by intraperitoneal injection of avertin (2%, v/v, 100 μl/10 g body weight) before imaging.

#### Construction of a mouse model of subcutaneous tumor

Mice were first allowed to acclimatize to the time laboratory culture environment for 1-2 weeks. Prepare the CT-26 cells to be transplanted, mix them 1:1 with stromal gel before injection, and then aspirate them with a 1 mL syringe. Sterilize the groin of mice with 75% alcohol, hold the syringe with the mixture of tumor cells and matrix gel in the right hand, enter the needle at an oblique angle of 45 degrees, be careful not to break through the peritoneum, inject the tumor cells mixed with matrix gel into the subcutaneous (the amount of tumor cells is about $1 \times 10^6$/mouse), withdraw the needle quickly, and then gently press the needle hole with the forefinger of the left hand for about 1 minute, and then return the mice to the feeding cage, and then observe the mice whether they awaken or not in 2-3 h. The growth of the tumor was observed in two weeks, and the mice were observed to grow. Tumor growth was observed within two weeks, and the next experiment could be carried out when the tumor volume was 1 cm × 1 cm.

#### Immunocytotherapy noninvasive real-time dynamic NIR-II-L ratiometric imaging

The C8R-DSNP nanosensor (200 μL, 200 μM) was first injected tail vein into the tumor site of Balb/c nude mice. Through the EPR effect of the nanoparticles, NK-92 cells ($1 \times 10^6$) were injected into the tail vein of mice 4 h later. Before and after tail vein injection, anaesthetised Balb/c nude mice were placed on an animal placement rack under laser. Real-time dynamic ratiometric imaging of the tumor site was performed on a commercialized NIR-II-L fluorescence imaging system (Shanghai United-Digital Biotech Co., Ltd.) by excitation with 980 nm and 808 nm lasers. Fluorescence images were collected with 1300 LP and 1400 LP.

#### Microscopic non-invasive NIR-II-L imaging of tumor

NK-92 cells were labelled in vitro by co-incubating DSNP (500 μL 200 μM) with NK-92 ($1\times10^6$) for 2 h. The supernatant was then incubated with DSNP (500 μL 200 μM) for 2 h. Afterwards, excess DSNP was removed from the supernatant by centrifugation (1000 rpm, 3 min) and washed once with 0.9% saline, the supernatant was removed by centrifugation again and the settled cells were resuspended in 0.9% saline. The labelled NK-92 cells were injected tail vein into Balb/c nude mice. Vessels were lit by tail vein injection of ICG. The NK-92 cells and tumor vessels were excited by a 980 nm laser and a 760 nm laser, respectively. Real-time kinetic NIR-II-L imaging of NK-92 cell behavior at the tumor site was performed using NIR-II-L fallout fluorescence microscopy.

#### Fluorescence imaging analysis of urinary metabolism in mice

Healthy mice, tumor mice, tumor mice injected with C8R-DSNP only, and mice injected with C8R-DSNP and NK-92 cells were placed into special metabolic cages. The urine and faeces of the mice were separated by urine and faeces separating funnel at the bottom of the cage, and the urine was collected from different groups and different time points (0 h, 24 h, 48 h), and the signals of ICG were collected by CCD imaging under the excitation of a 760 nm exciter.

#### Apoptosis detection of tissues

**TUNEL staining:** Tumor tissue sections were washed twice with xylene for 5 min each, and once with graded ethanol (100%, 95%, 90%, 80%, 70%) for 3 min each, followed by rinsing twice with PBS, gently discarding the PBS and blotting the excess liquid with filter paper. Tissues were treated with Proteinase K working solution for 15-30 min at 21-37 °C or with cell permeabilisation solution for 8 min, followed by 2 rinses in PBS. Prepare TUNEL reaction mix by mixing 50 μL TdT + 450 μL fluorescein-labelled dUTP solution, and react for 10-30 min at room temperature to 37°C. After the slides are dried, add 50 μL TUNEL reaction mix to the specimens, cover slips or sealing film in a dark humidor and react for 60 min at 37 °C, followed by rinsing 3 times with PBS. Add the anti-fluorescence quenching sealing solution (HY-K1042) to seal the film and observe the fluorescence effect under fluorescence microscope.

**Ki67 immunohistochemical staining:** Tumor sections were first treated with antigenic thermal repair. The repaired sections were washed with tap water to clean the sections, after which they were placed in a vat containing 3% horseradish peroxidase (HRP) staining vat for 10 min, and then removed and the sections were washed with tap water. The water left on the sections was removed and the area of tissue to be tested on the slide was circled with an immunohistochemical pen and placed in PBST buffer. After removing the PBST buffer, 3 drops of Ki-67 antibody reagent were added to each section (depending on the size of the section to completely cover the sectioned tissue) and incubated for 30 min at room temperature. After the sections were washed in PBST buffer, 100 μL of enzyme-labelled goat anti-mouse IgG polymer was added to each section to completely cover the sectioned tissue and incubated for 30 min at room temperature. Color development was performed with freshly prepared diaminobenzidine (DAB) chromogenic solution, and about 100 μL of DAB chromogenic solution was added to each section to completely cover the section tissues for 1-5 min. Sections were washed with tap water and re-stained with hematoxylin staining solution, then dehydrated, transparent and sealed. Finally, DAB was catalyzed by HRP to form brown deposits at the antigenic site. The presence of the target antigen and its expression was determined by observing the brown areas under a light microscope.

#### Bio-safety Testing

**Immunohistochemistry (IHC):** (1) The heart, liver, spleen, lungs and kidneys of each group of mice were dissected, washed in PBS and fixed in 4% paraformaldehyde for 24 h. Afterwards, the tissues were removed from the fixative and rinsed under running water to remove residual fixative and crystals generated. (2) Tissue samples were processed through embedding, sectioning, dewaxing and hydration, antigen repair, inactivation, sealing, primary antibody incubation, secondary antibody incubation, colour development, re-staining and sealing (this step was done by Shanghai yanyue Biotechnology Co., Ltd.), and then finally, the samples were subjected to immunohistochemistry under the microscope.

**Blood routine examination:** Blood is taken from the orbital venous plexus (sinus) of the mouse by anesthetizing the animal, placing it in a lateral position on a flat surface, and pressing the ends of the neck with the forefinger and thumb to make venous blood return to the head difficult and the eyeball protruding. A rigid capillary glass tube, at an angle of 45° to the face of the mouse, was punctured between the eyelid and the eyeball from the inner corner of the eye, moving towards the base of the eye with a slight rotation to incise the venous plexus. The mice were punctured to a depth of approximately 2 to 3 mm, and the extraction vessel was pushed and withdrawn at the same time. Once the desired amount of blood was taken, the pressure applied to the neck was removed and the blood collector was pulled out to collect the blood.

#### Statistics and reproducibility

All images were processed using MATLAB 2022 and ImageJ FIJI v4.3.0 software. Graphs were plotted using GraphPad Prism 10 and statistically analysand as appropriate values are shown as mean ± s.e.m.; $*P < 0.05$, $**P < 0.01$, $***P < 0.001$, $****P < 0.0001$. All experiments were performed independently at least 3 times.

#### Results and Discussion

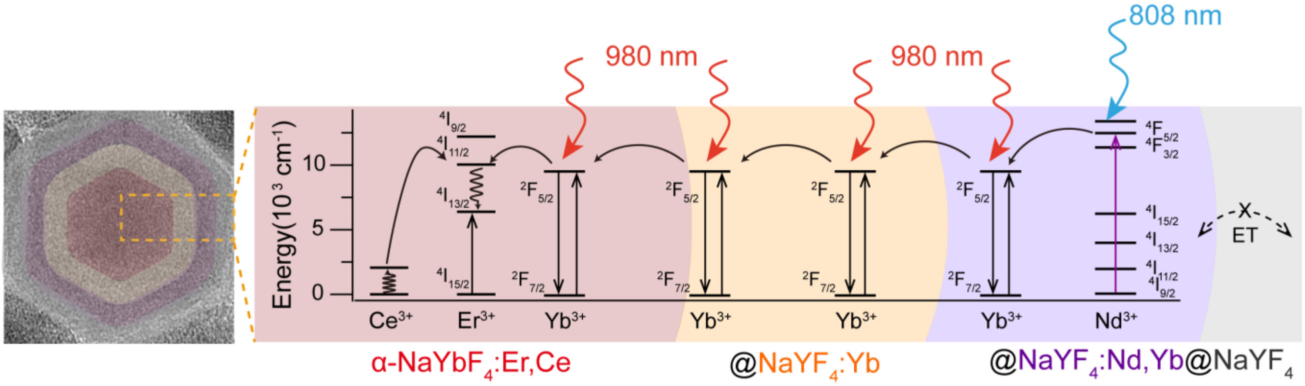

**Figure S1.** Luminescence mechanism of DSNP and its structure. Energy transfer mechanism of NIR-II-L fluorescence in α-$NaYbF_4$:Er,Ce@$NaYF_4$:Yb@$NaYF_4$:Nd,Yb@$NaYF_4$ (DSNP).

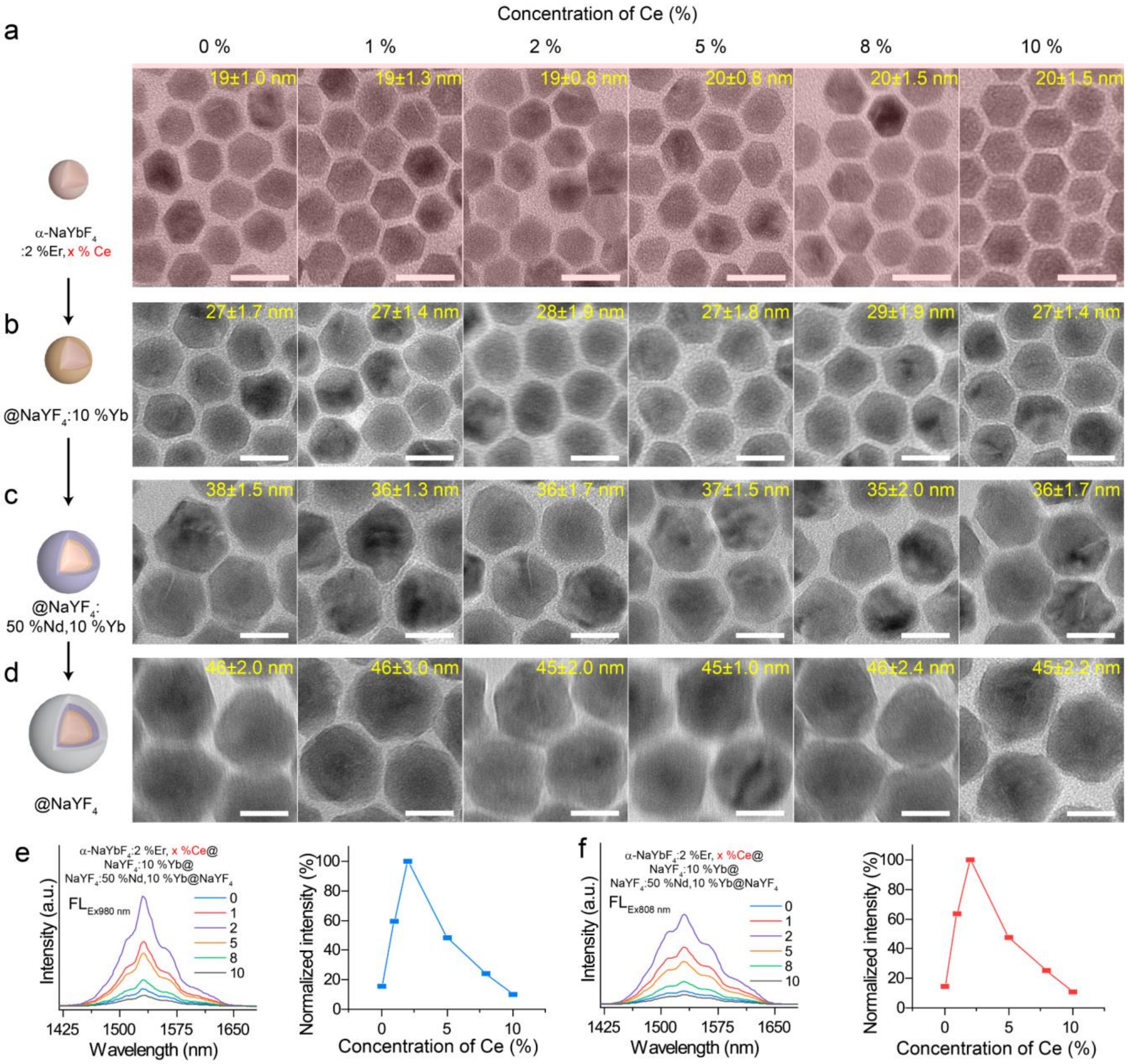

**Figure S2. Characterization of core-shell-shell-shell DSNPs with different $Ce^{3+}$ contents.** (**a**)-(**d**) TEM images of α-$NaYbF_4$:Er,x%Ce@$NaYF_4$:Yb@$NaYF_4$:Nd,Yb @$NaYF_4$ nanoparticles with $Ce^{3+}$ contents ranging from 0 mmol% to 10 mmol%. NIR-II-L emission spectra of nanoparticles of α-$NaYbF_4$:Er,x%Ce@$NaYF_4$:Yb@$NaYF_4$:Nd,Yb@$NaYF_4$ under (**e**) $FL_{Ex980\ nm}$ excitation or (**f**) $FL_{Ex808\ nm}$ excitation, and their normalized statistical plots. Scale bar, 25 nm.

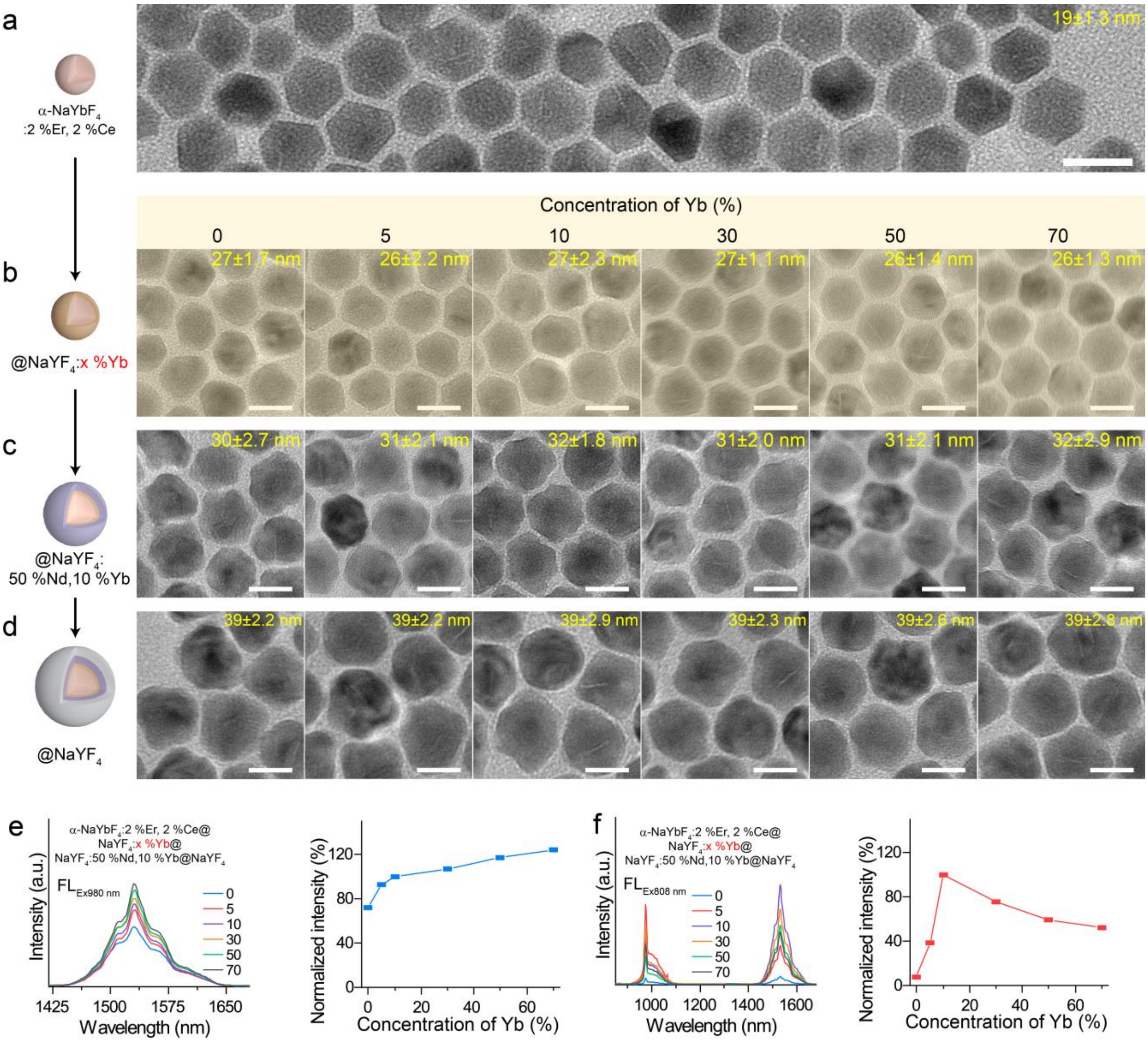

**Figure S3. Characterization of core-shell-shell-shell DSNPs with different $Yb^{3+}$ contents in $NaYF_4$: Yb shell.** (**a**)-(**d**), TEM images of α-$NaYbF_4$:Er,2%Ce@$NaYF_4$:x%Yb@$NaYF_4$:Nd,Yb@$NaYF_4$ nanoparticles with $Yb^{3+}$ contents ranging from 0 mmol% to 70 mmol%. NIR-II-L emission spectra of nanoparticles of $NaYbF_4$:Er,2%Ce@$NaYF_4$: x%Yb shell@$NaYF_4$:Nd,Yb@$NaYF_4$ under (**e**) $FL_{Ex980\ nm}$ excitation or (**f**) $FL_{Ex808\ nm}$ excitation and their normalized statistical plots. Scale bar, 25 nm.

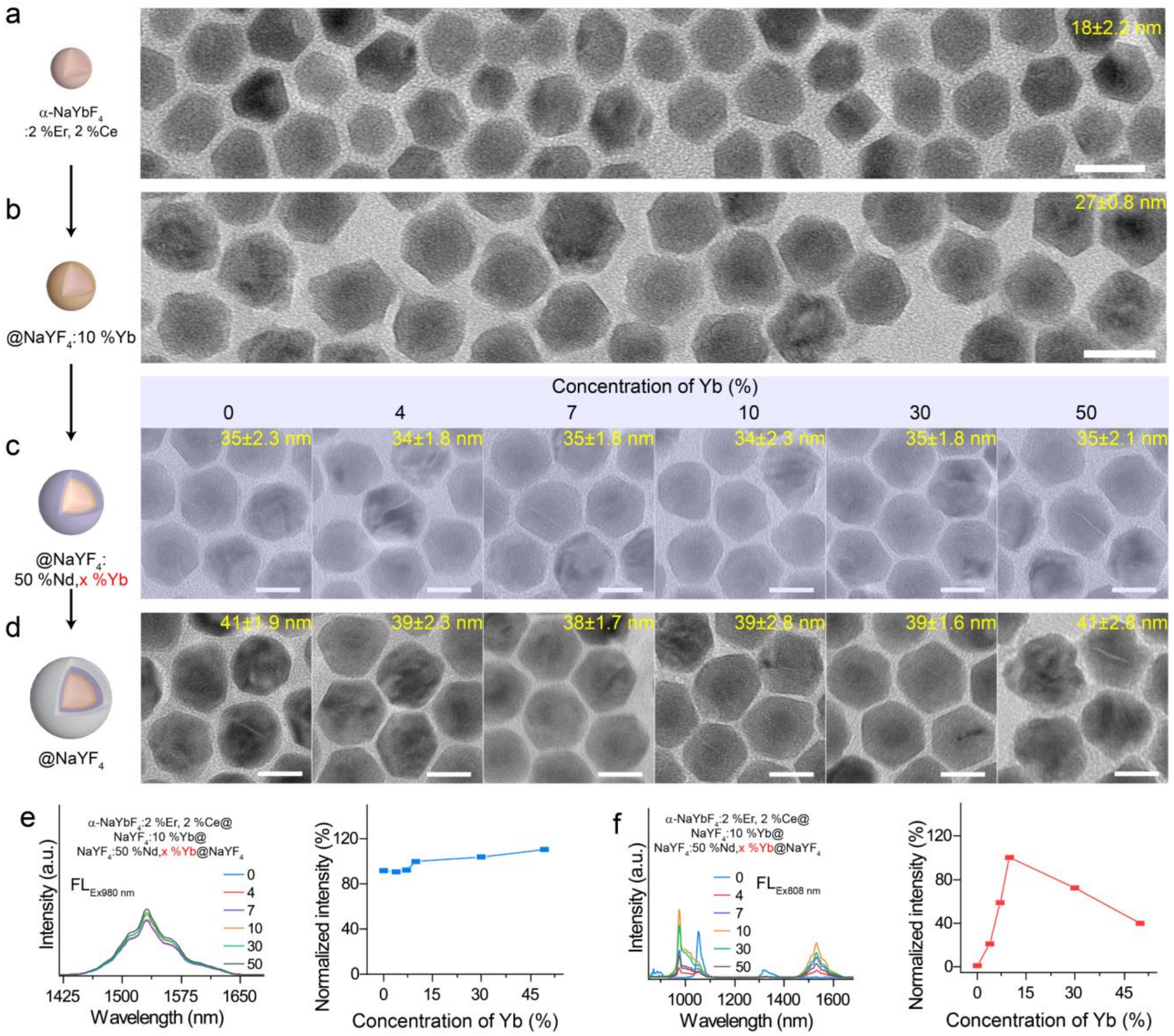

**Figure S4. Characterization of core-shell-shell-shell DSNPs with different $Yb^{3+}$ contents in $NaYF_4$: Nd, Yb shell.** (**a**)-(**d**), TEM images of α-$NaYbF_4$:Er,2%Ce@$NaYF_4$:10%Yb@$NaYF_4$:Nd,x%Yb@$NaYF_4$ nanoparticles with $Yb^{3+}$ contents ranging from 0 mmol% to 50 mmol%. NIR-II-L emission spectra of nanoparticles of $NaYbF_4$:Er,2%Ce@$NaYF_4$:10%Yb@$NaYF_4$:Nd,x%Yb@$NaYF_4$ under (**e**) $FL_{Ex980\ nm}$ excitation or (**f**) $FL_{Ex808\ nm}$ excitation and their normalized statistical plots. Scale bar, 25 nm.

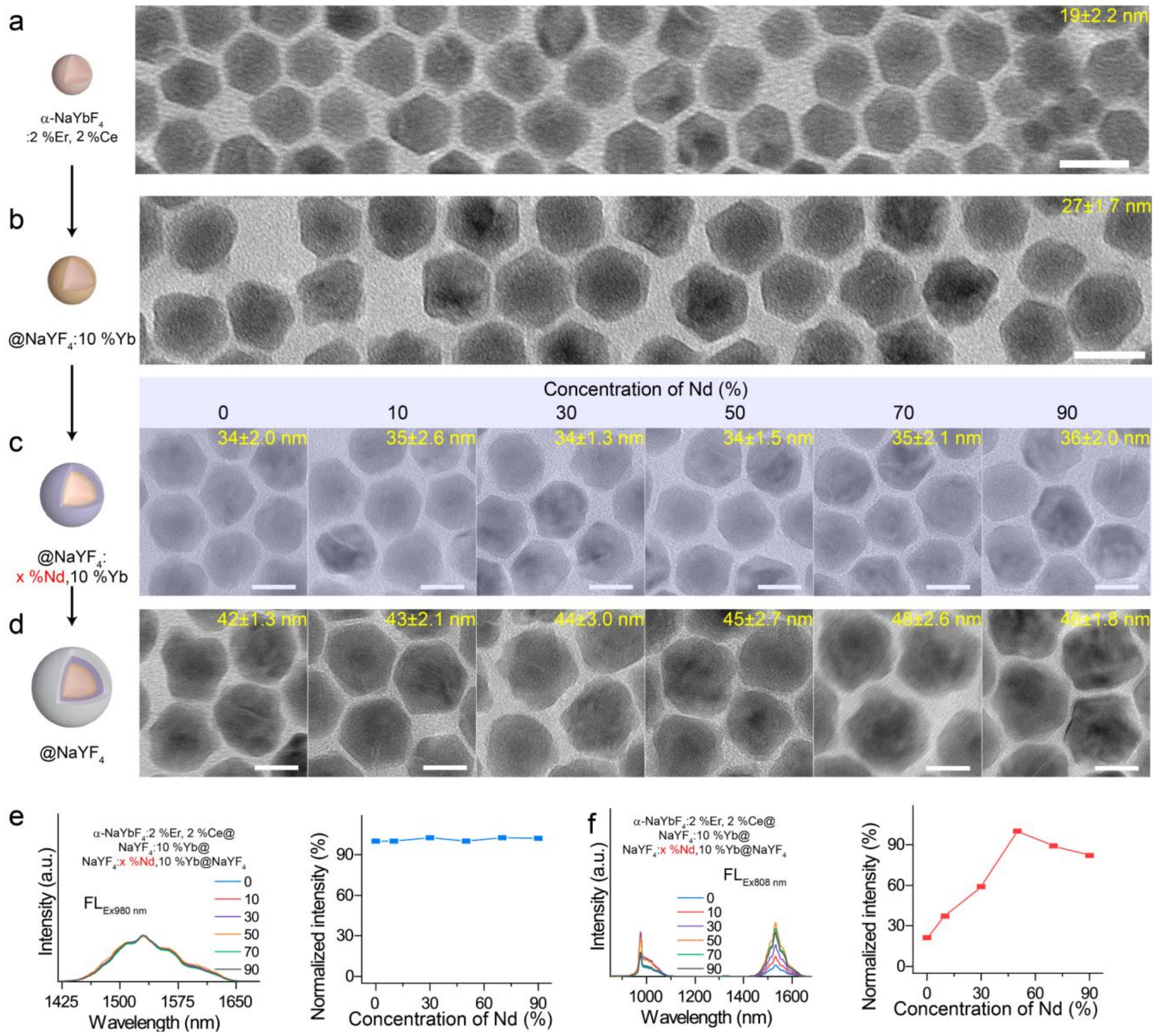

**Figure S5. Characterization of core-shell-shell-shell DSNPs with different $Nd^{3+}$ contents.** (**a**)-(**d**), TEM images of α-$NaYbF_4$:Er,2%Ce@$NaYF_4$:10%Yb@$NaYF_4$:x%Nd,10%Yb@$NaYF_4$ nanoparticles with $Nd^{3+}$ contents ranging from 0 mmol% to 90 mmol%. NIR-II-L emission spectra of nanoparticles of $NaYbF_4$:Er,2%Ce@$NaYF_4$:10%Yb@$NaYF_4$:x%Nd,10%Yb@$NaYF_4$ under (**e**) $FL_{Ex980\ nm}$ excitation or (**f**) $FL_{Ex808\ nm}$ excitation and their normalized statistical plots. Scale bar, 25 nm.

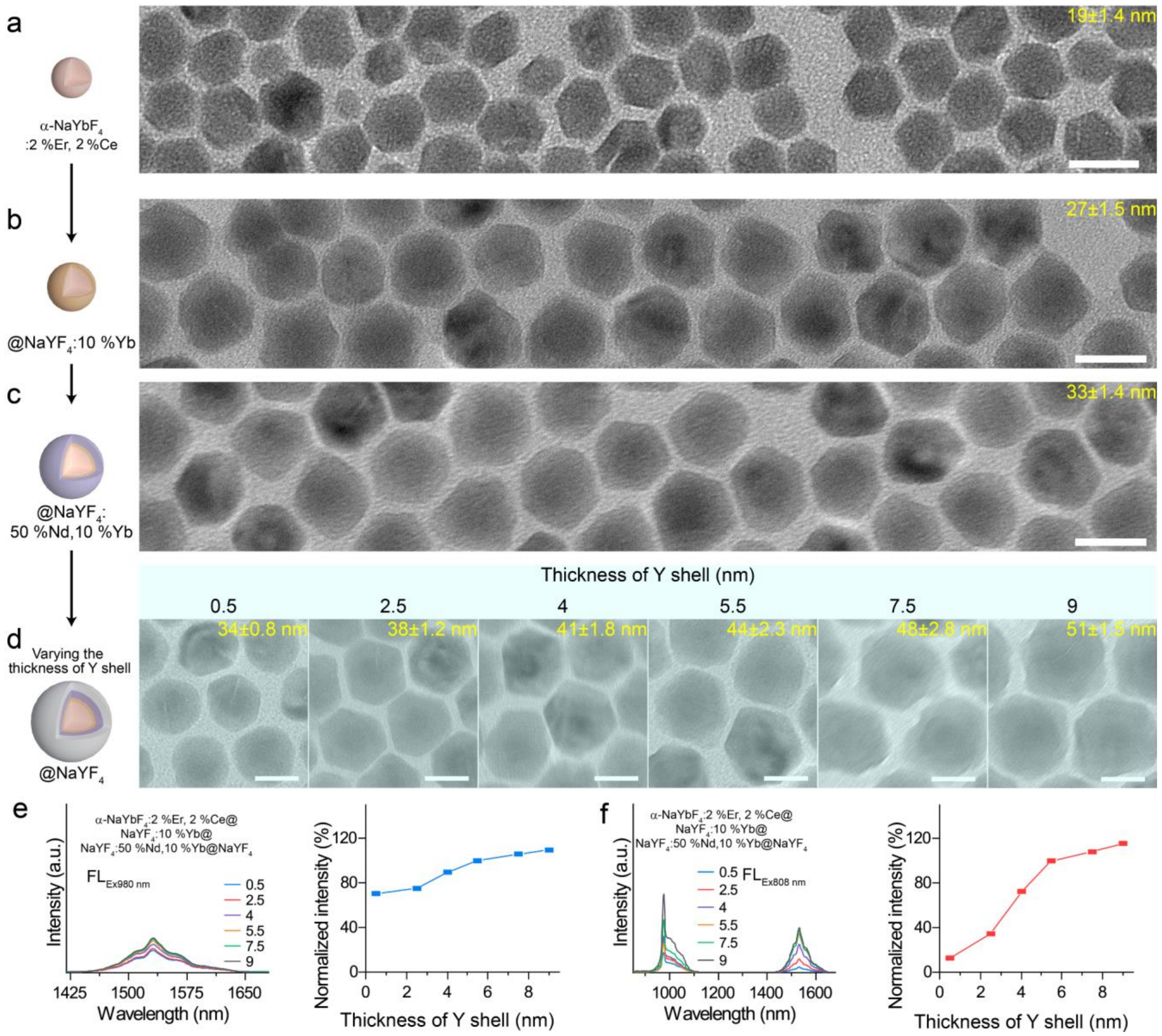

**Figure S6. Characterization of core-shell-shell-shell DSNPs with different thickness of Y shell.** (**a**)-(**d**), TEM images of α-$NaYbF_4$:Er,2%Ce@$NaYF_4$:10%Yb@$NaYF_4$:50%Nd,10%Yb@$NaYF_4$ nanoparticles with $NaYF_4$ various thicknesses from 0.5 nm to 9 nm. NIR-II-L emission spectra of nanoparticles of $NaYbF_4$:Er,2%Ce@$NaYF_4$:10%Yb@$NaYF_4$:50%Nd,10%Yb@$NaYF_4$ under (**e**) $FL_{Ex980\ nm}$ excitation or (**f**) $FL_{Ex808\ nm}$ excitation and their normalized statistical plots. Scale bar, 25 nm.

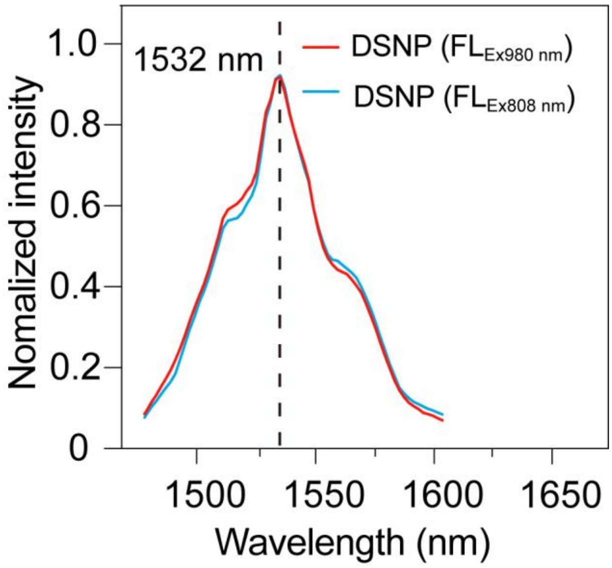

**Figure S7.** $FL_{Ex980\ nm}$ and $FL_{Ex808\ nm}$ of water-soluble DSNP shows consistent NIR-II-L emission spectral shape.

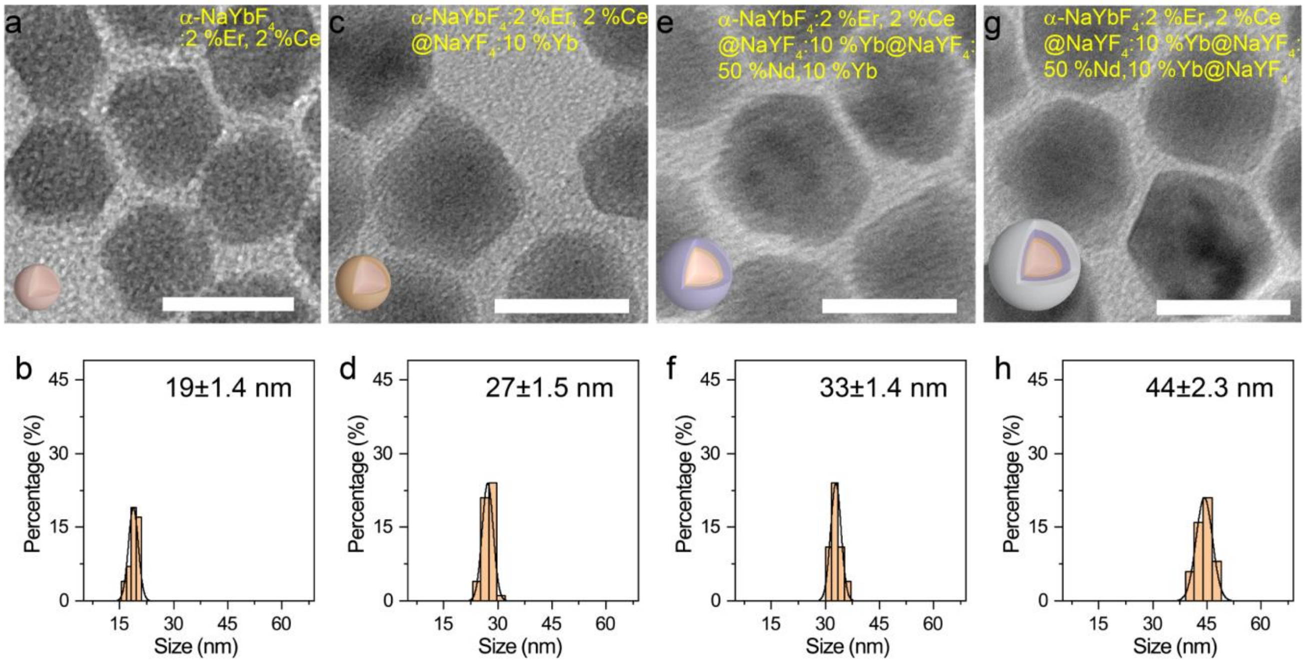

**Figure S8.** Electron transmission microscopy (TEM) images of core-shell-shell-shell structured nanoparticles after stepwise encapsulation (a, c, e and g) and particle size statistics (b, d, f and h). Scale bar, 50 nm.

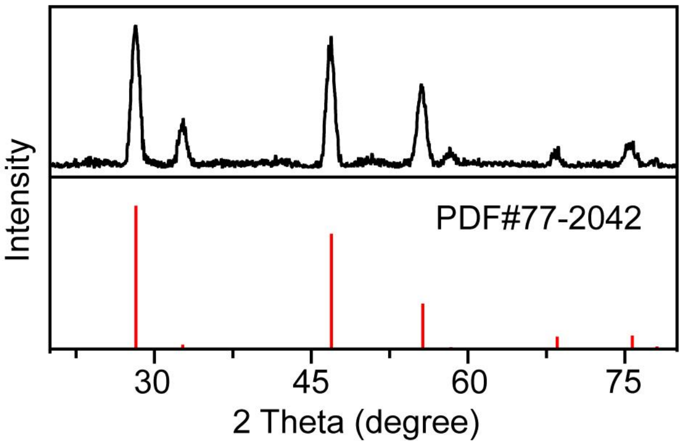

**Figure S9.** X-ray diffraction (XRD) pattern of C8R-DSNP, showing consistent structure with peaks corresponding to those of pure α-$NaYF_4$ crystals.

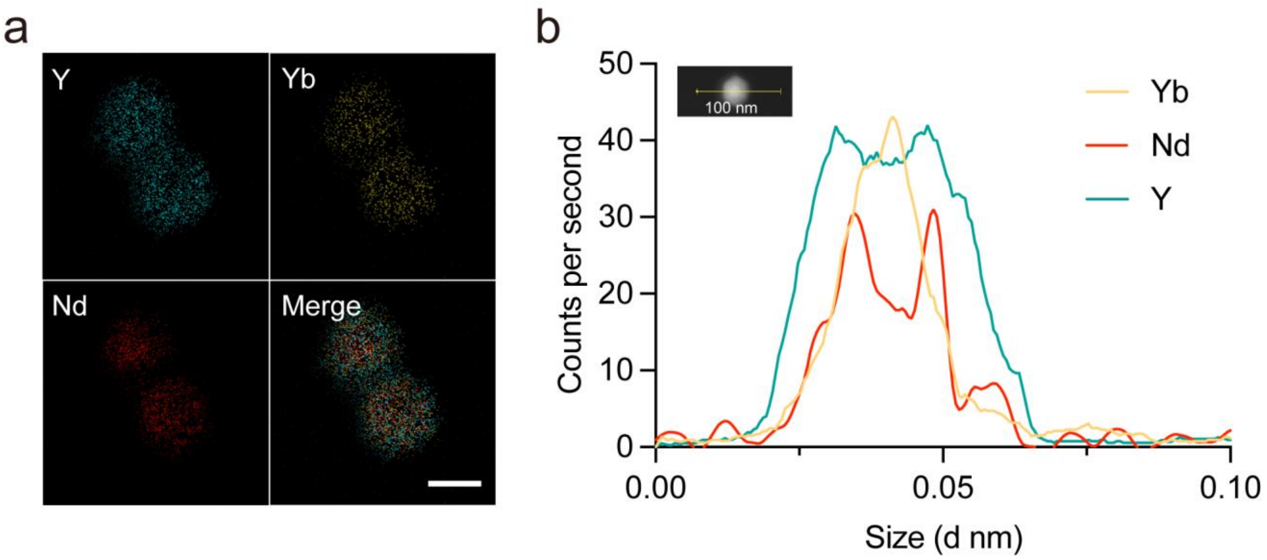

**Figure S10.** (a) **Elemental mapping analysis images of DSNP (scale bar, 25 nm).** (b) TEM line scanning was used to determine the relative content of elements in different regions of the DSNP particles, scale bar, 100 nm.

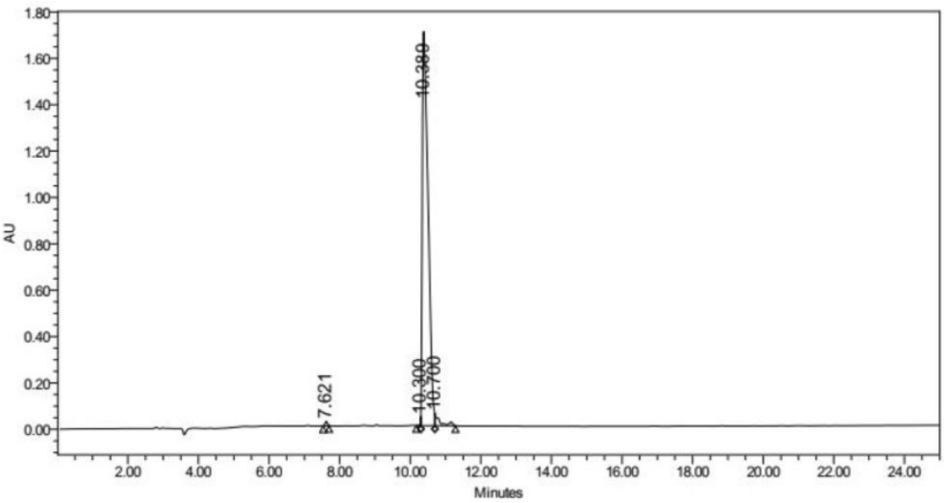

| | RT | Area | % Area | Height |
|---|---|---|---|---|
| 1 | 7.621 | 83056 | 0.42 | 18244 |
| 2 | 10.300 | 27078 | 0.14 | 37417 |
| 3 | 10.389 | 19332131 | 96.99 | 1713152 |
| 4 | 10.700 | 489645 | 2.46 | 50798 |

**Figure S11.** High-performance liquid chromatography (HPLC) analysis shows the synthesized caspase-8-specific substrate IETDC with a purity of 96.99%.

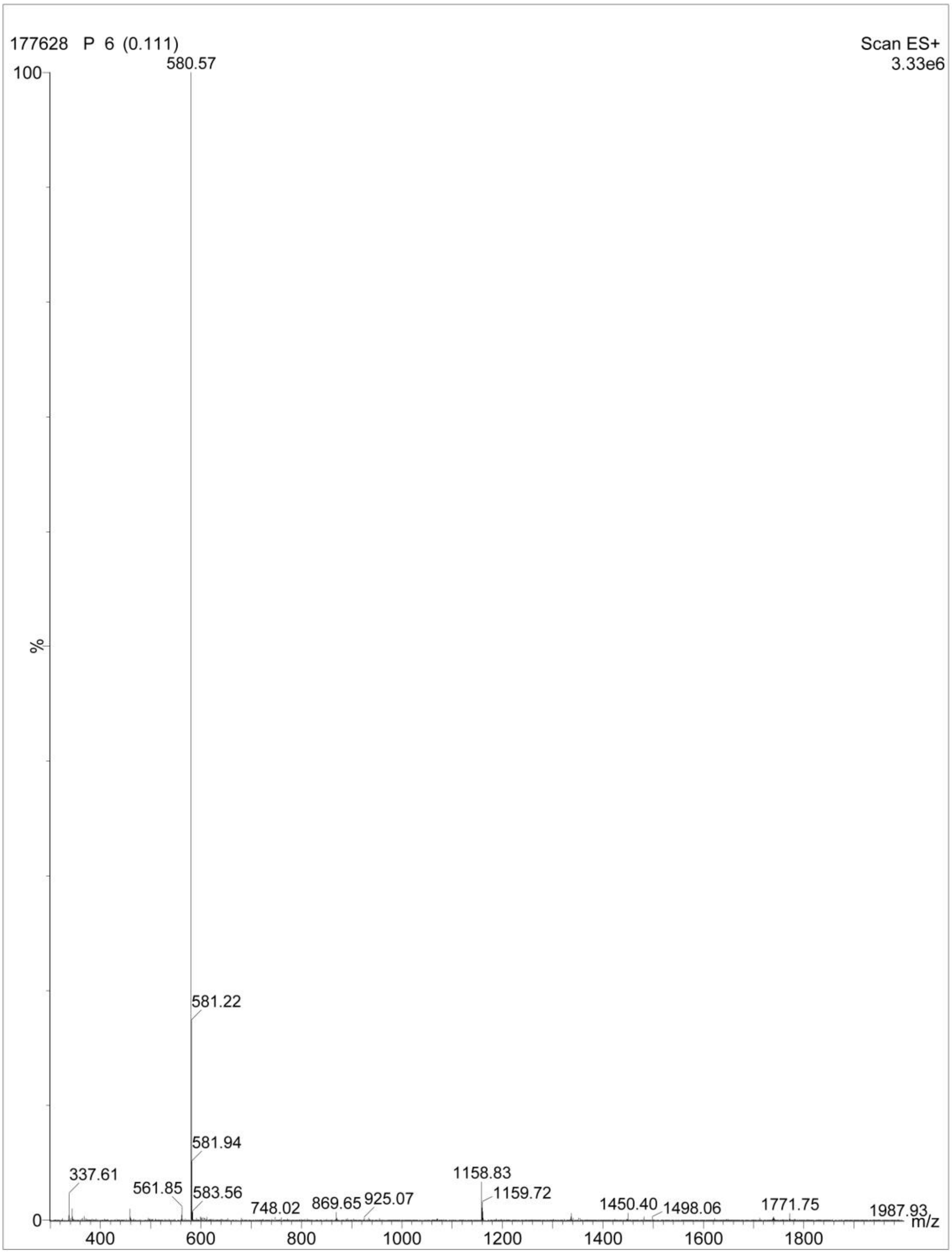

**Figure S12.** High-resolution mass spectrometry of the caspase-8-specific substrate IETDC. The molecular weight of the compounds was deduced from the m/z value of the highest intensity peak in agreement with IETDC (MW: 579.62).

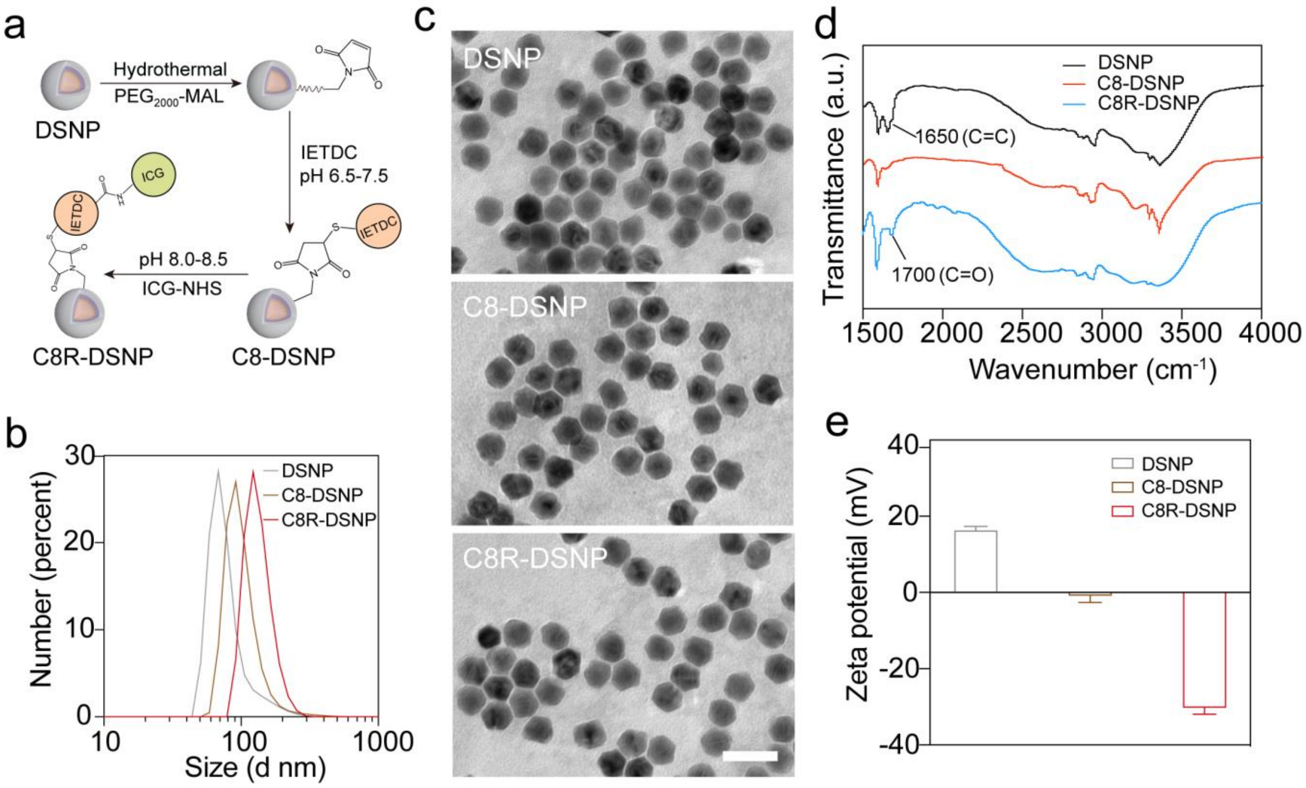

**Figure S13. Characterization of the DSNP modification for biological application.** (a) Schematic representation of the two-step modification of nanoparticles and (b) hydrodynamic nanoparticle size analysis (right). (c) TEM before and after the two-step conjugation reaction. a.u., arbitrary unit. scale bar, 50 nm. (d) FTIR spectra of DSNP, C8-DSNP and C8R-DSNP. (e) Zeta potential of DSNP, C8-DSNP and C8R-DSNP.

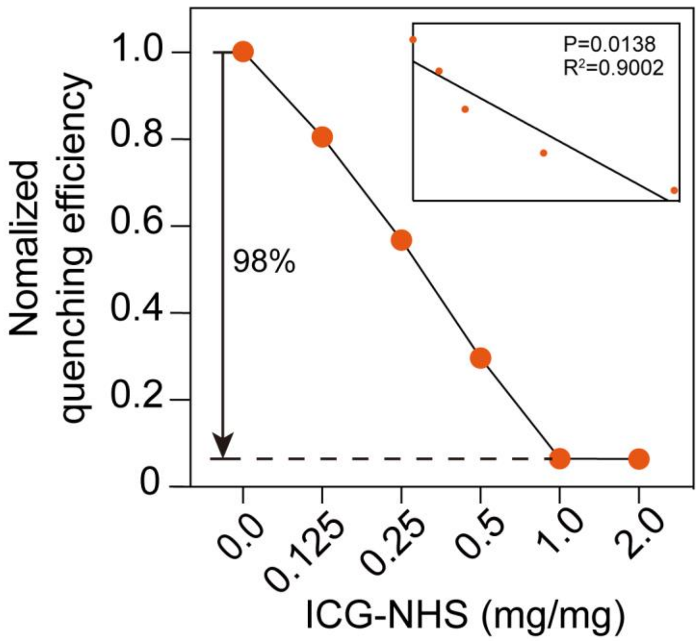

**Figure S14.** Relationship for NIR-II-L ratiometric fluorescence ($FL_{Ex808\ nm}$/ $FL_{Ex980\ nm}$) over ICG-NHS concentrations. Inset shows the linear relationship.

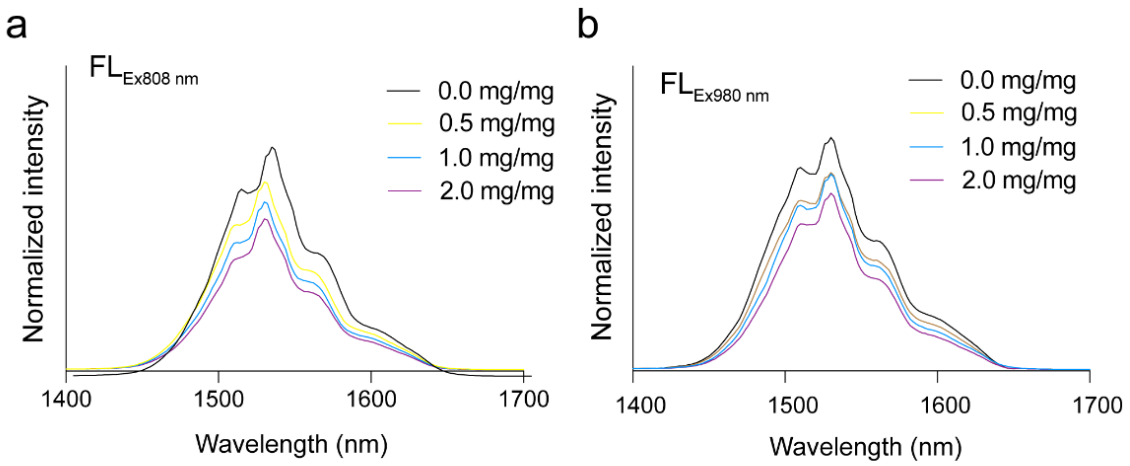

**Figure S15 $FL_{Ex808\ nm}$ and $FL_{Ex980\ nm}$ of DSNP mixed with different concentrion of ICG in trichloromethane solution.** Emission spectra at 1532 nm under 808 nm excitation (**a**) or 980 nm excitation (**b**). Physical mixing of DSNP with ICG had no effect on the NIR-II-L fluorescence of DSNP under 808 nm and 980 nm excitation.

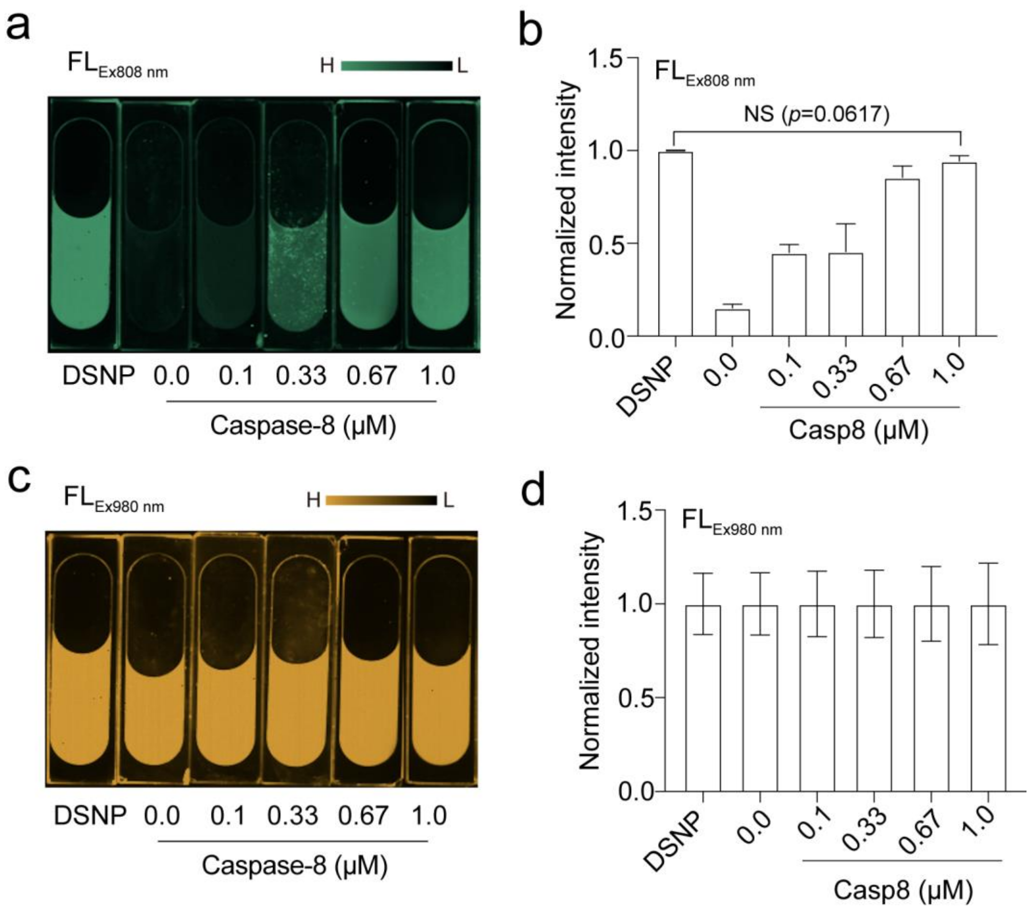

**Figure S16. Comparison of NIR-II-L imaging with different caspase-8 concentration under 808 nm or 980 nm.** (**a**) NIR-II-L fluorescence images of C8R-DSNP after the addition of different concentrations of caspase-8 under 808 nm. (**b**) Single-channel ($FL_{Ex808\ nm}$) statistical analyses of fluorescence intensities. (**c**) NIR-II-L fluorescence images of C8R-DSNP after the addition of different concentrations of caspase-8 under 980 nm. (**d**) Single-channel ($FL_{Ex980\ nm}$) statistical analyses of fluorescence intensities.

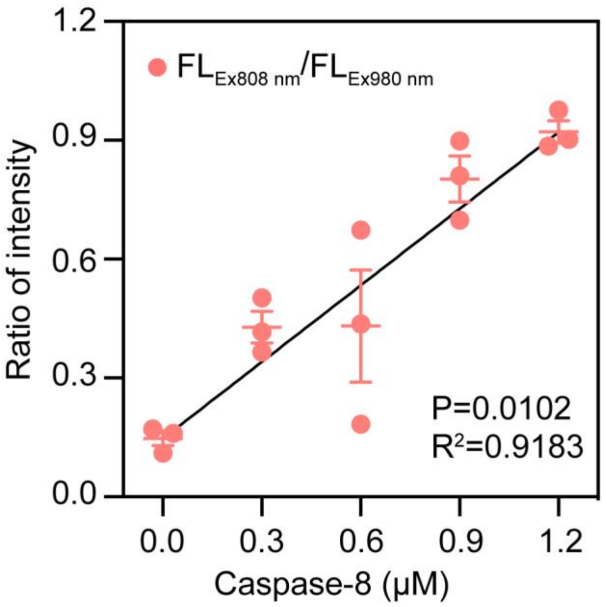

**Figure S17.** Linear analysis of caspase-8 concentration versus ratiometric fluorescence. Statistical values are expressed as mean ± s.e.m.; NS indicates no significance (n = 3).

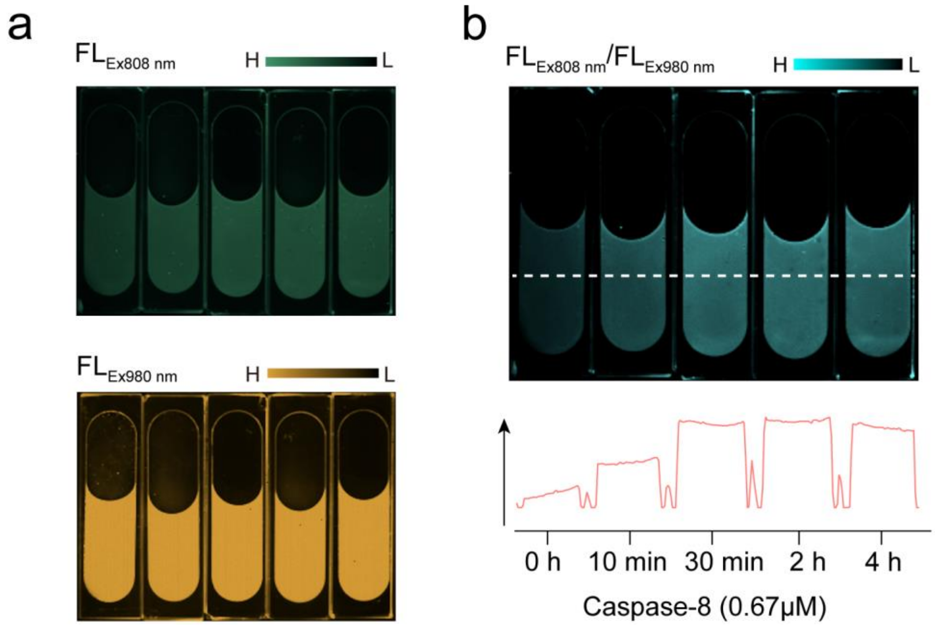

**Figure S18. NIR-II-L fluorescence imaging of C8R-DSNP after incubation with caspase-8 for various time.** (**a**) NIR-II-L images of C8R-DSNP excited at 808 nm and 980 nm by adding 0.67 μM caspase-8 for 0, 10, 30, 120 and 240 min. (**b**) Corresponding NIR-II-L ratiometric fluorescence images calculated from (**a**). H: high fluorescence intensity; L: low fluorescence intensity.

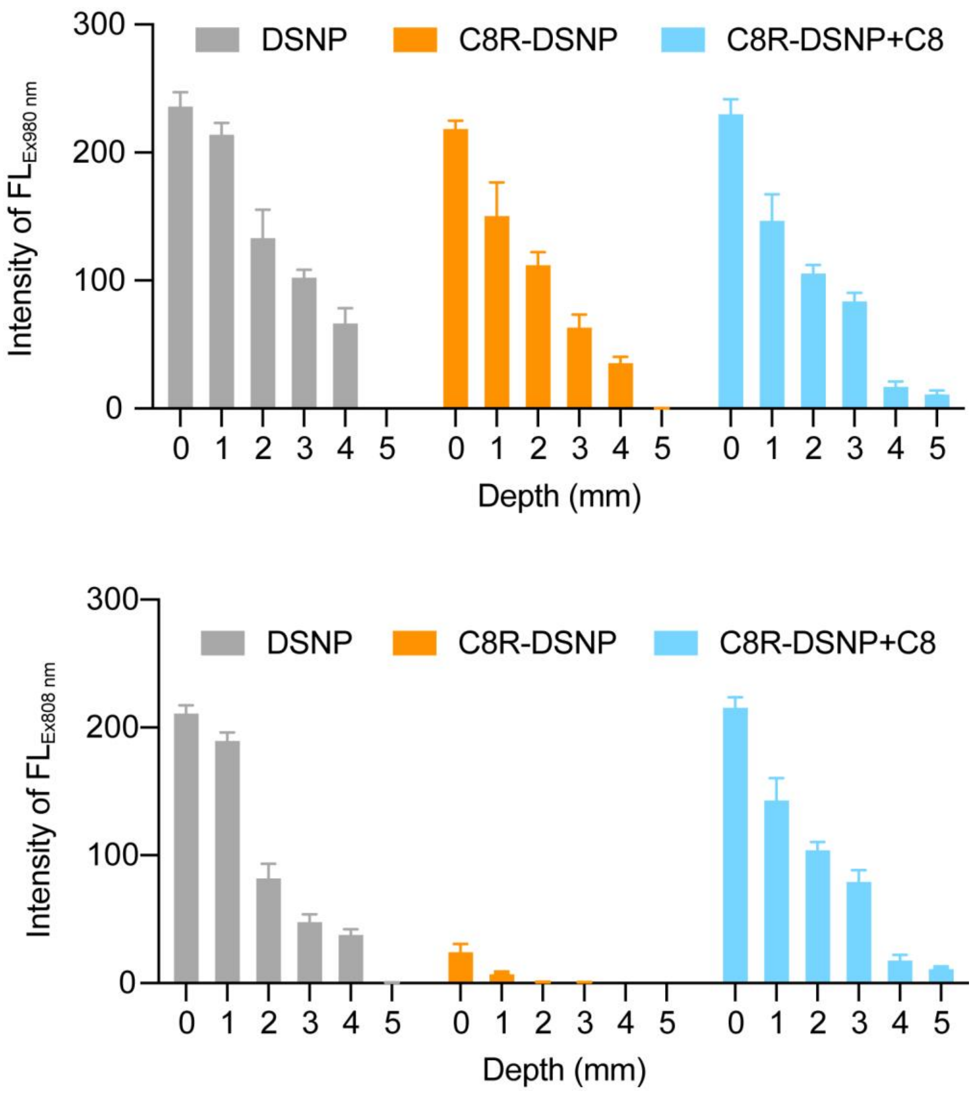

**Figure S19.** Statistics on fluorescence intensity of C8R-DSNP filled capillaries before and after caspase-8 response under various mimic tissue depth.

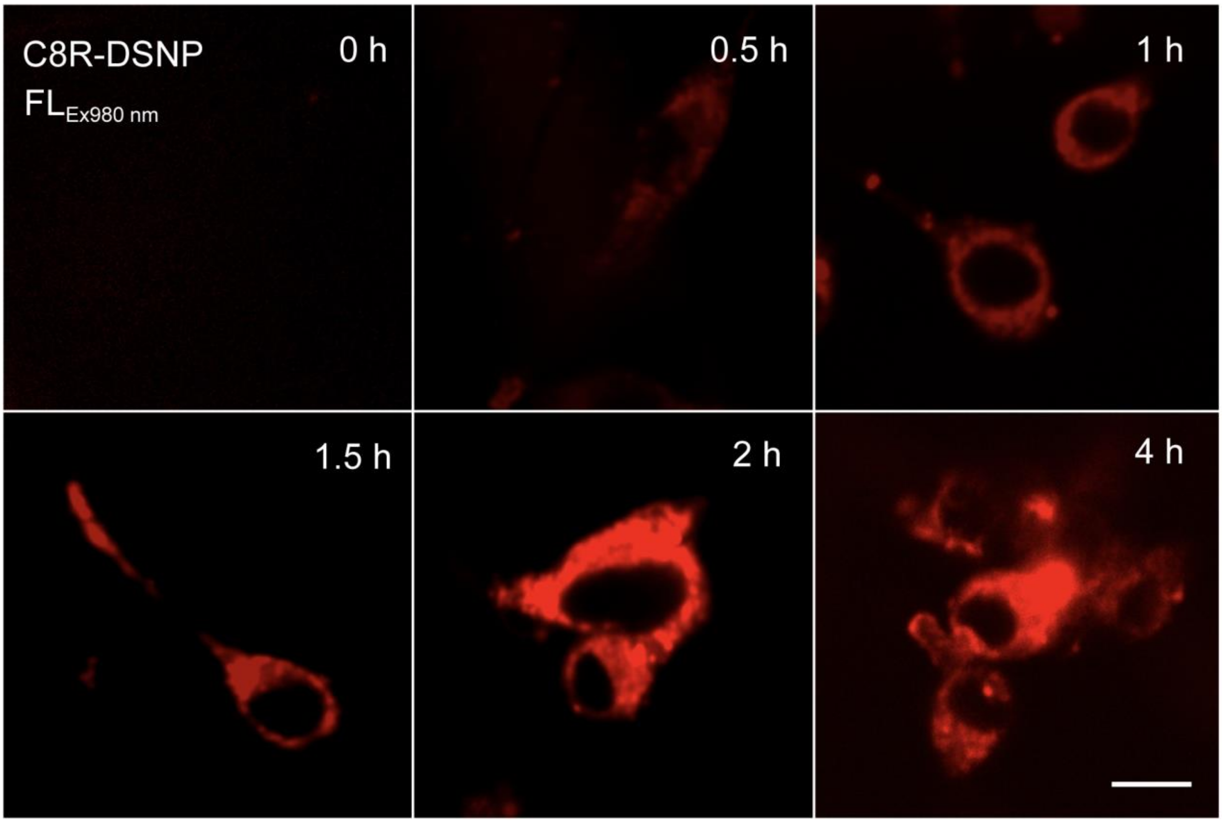

**Figure S20.** Endocytosis of C8R-DSNP by CT-26. Confocal microscopy imaging showed the best phagocytosis of C8R-DSNP by cells at 2 h.

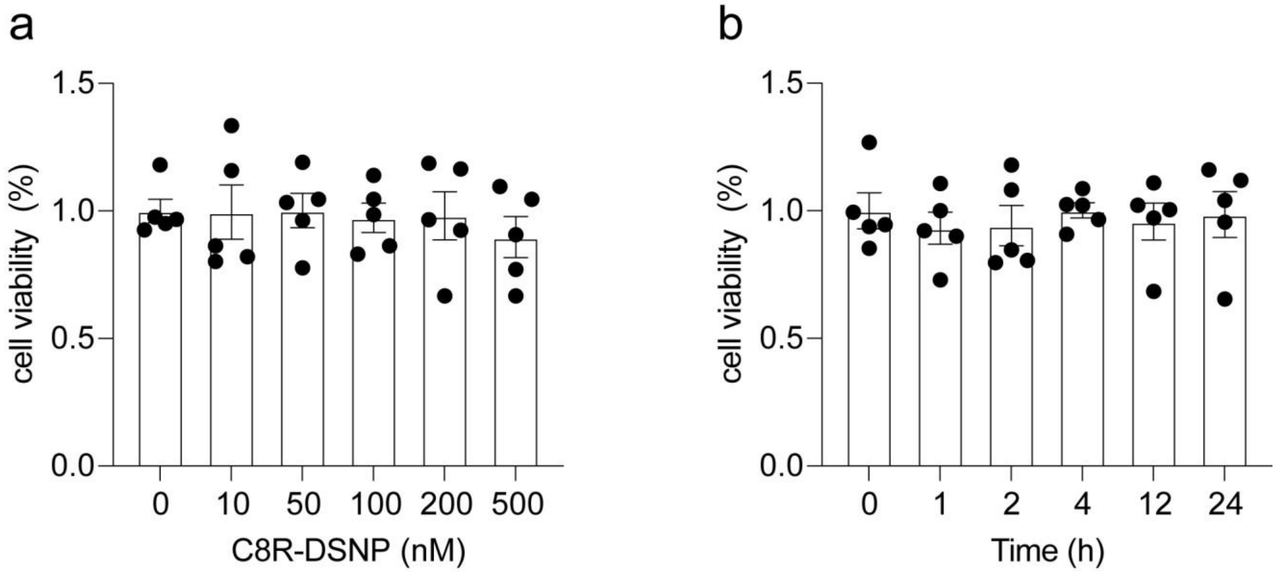

**Figure S21. The cytotoxicity assay of C8R-DSNP.** (**a**) C8R-DSNP showed no significant toxicity to cells up to 500 nM. (**b**) Cell viability with 500 nM C8R-DSNP for 24 h. The results show no significant inhibitory effect on cell proliferation. Statistical values are expressed as mean ± s.e.m. (n = 5).

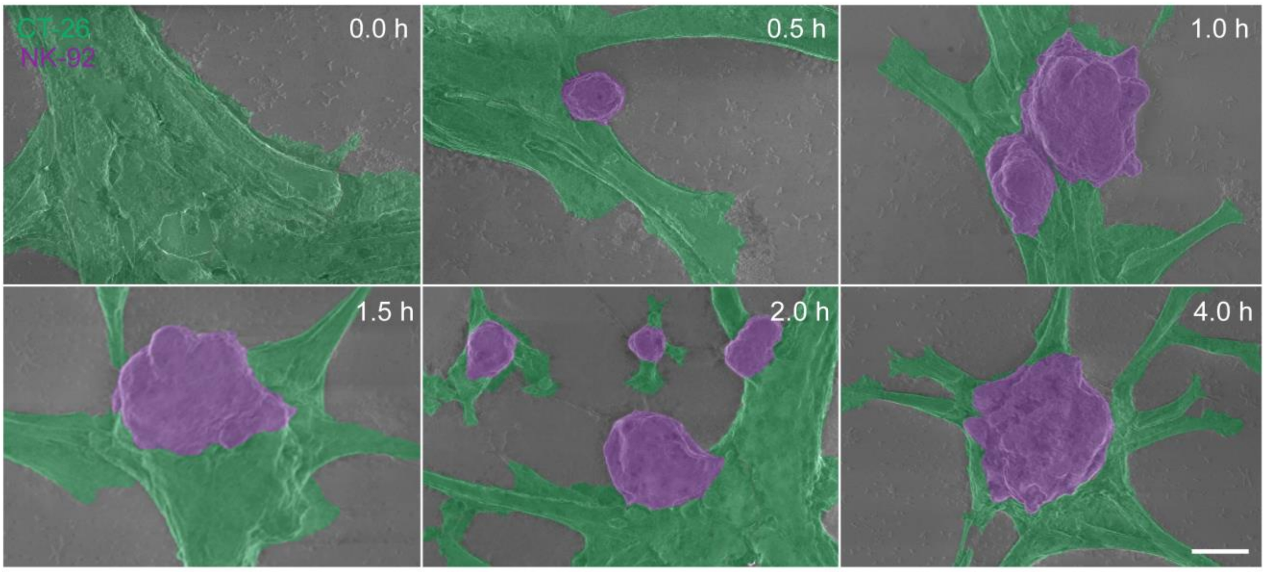

**Figure S22.** Representative scanning electron microscope image of NK-92 cells (purple) interacting with CT-26 cells (green), scale bar, 10 μm.

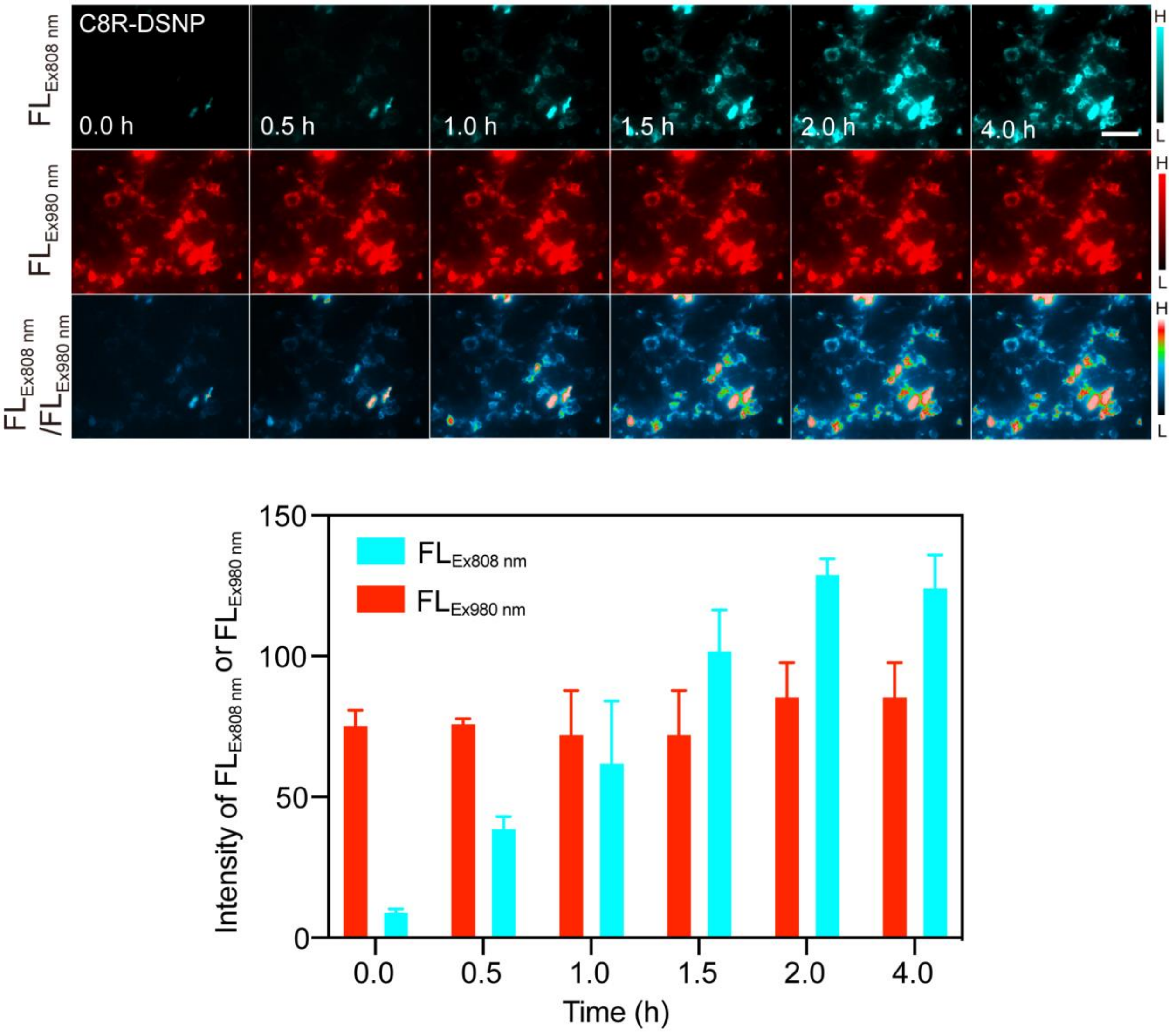

**Figure S23.** Corresponding NIR-II-L images of CT-26 cells under 980 nm and 808 nm excitation at different time points (0–4 h). NIR-II-L ratiometric fluorescence results were also shown**.** H: high fluorescence intensity; L: low fluorescence intensity. (**b**) Statistical analysis of $FL_{Ex980\ nm}$ and $FL_{EX808\ nm}$ at different time points**.** Statistical values are expressed as mean ± s.e.m. (n = 3).

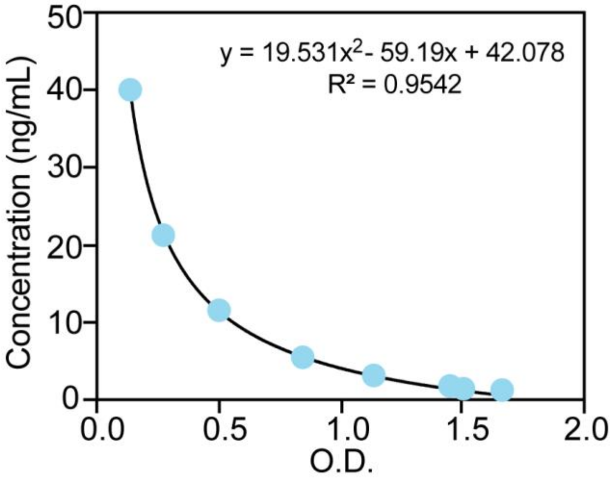

**Figure S24.** ELISA standard curve for caspase-8 protease concentration.

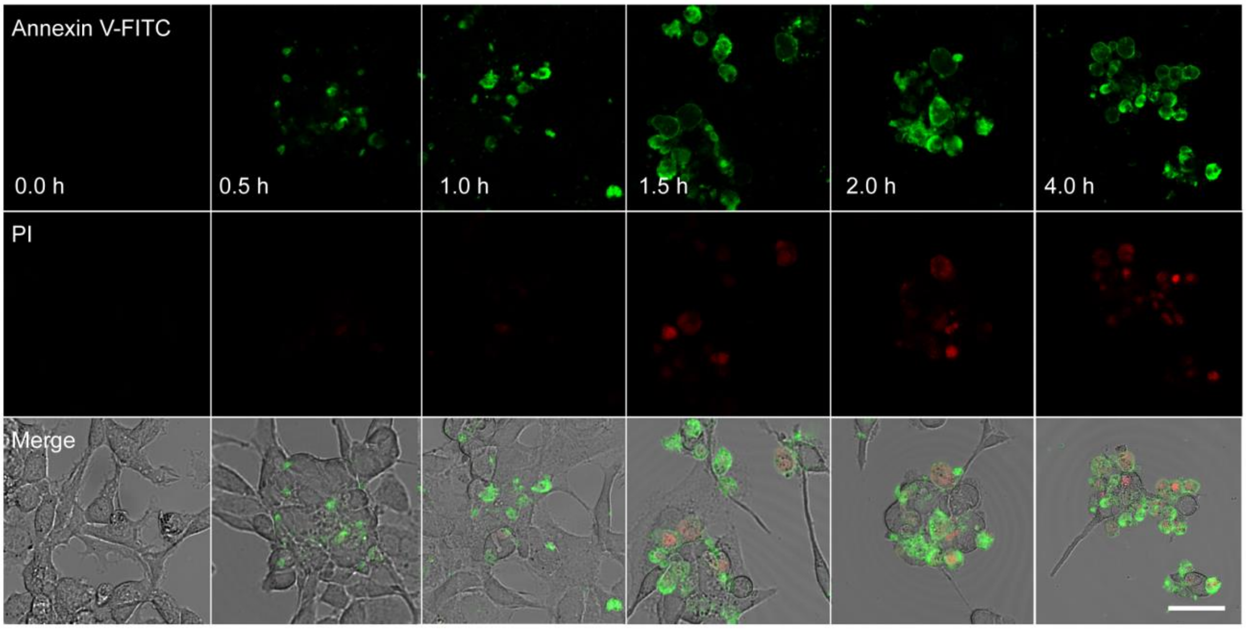

**Figure S25.** Confocal images of cell apoptosis (early apoptosis in green and late apoptosis in red) upon NK-92 cells and CT-26 cells interactions for different time.

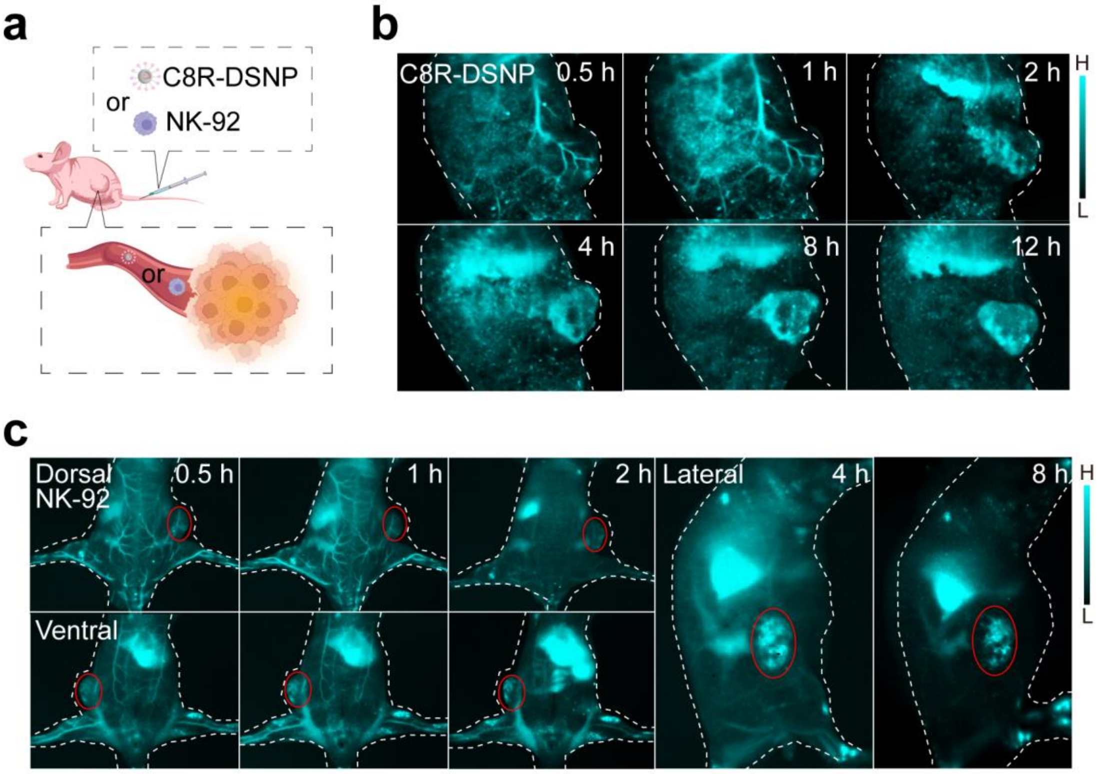

**Figure S26.** (**a**) Schematic illustration of in vivo circulation of injected C8R-DSNP or NK-92 cells in mice. In vivo NIR-II-L imaging of mice at different times after injection of C8R-DSNP (**b**) and NK-92 cells (**d**).

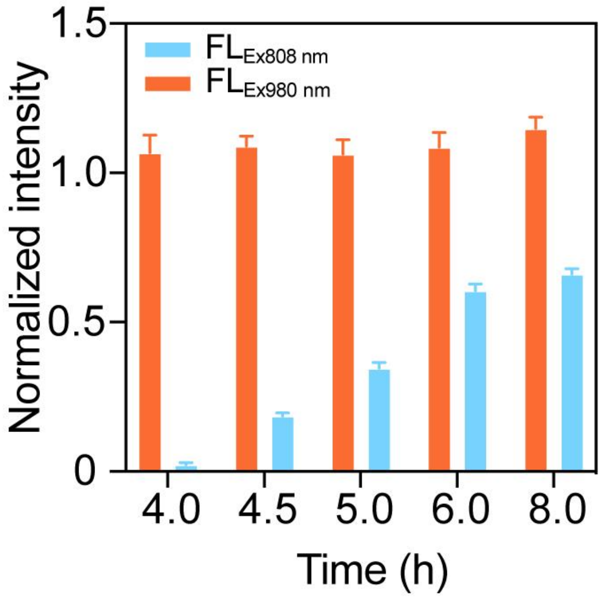

**Figure S27.** Statistical analysis of $FL_{Ex980\ nm}$ and $FL_{EX808\ nm}$ at different time points**.** Statistical values are expressed as mean ± s.e.m. (n = 3).

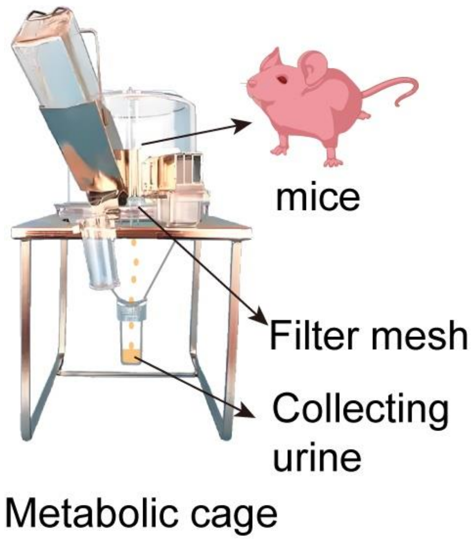

**Figure S28.** Schematic diagram of urinary metabolite collection in mice. Mice are monitored for their metabolic status by housing them in specialized metabolic cages. The excretion collection system includes components for collecting urine and feces, facilitating subsequent analysis.

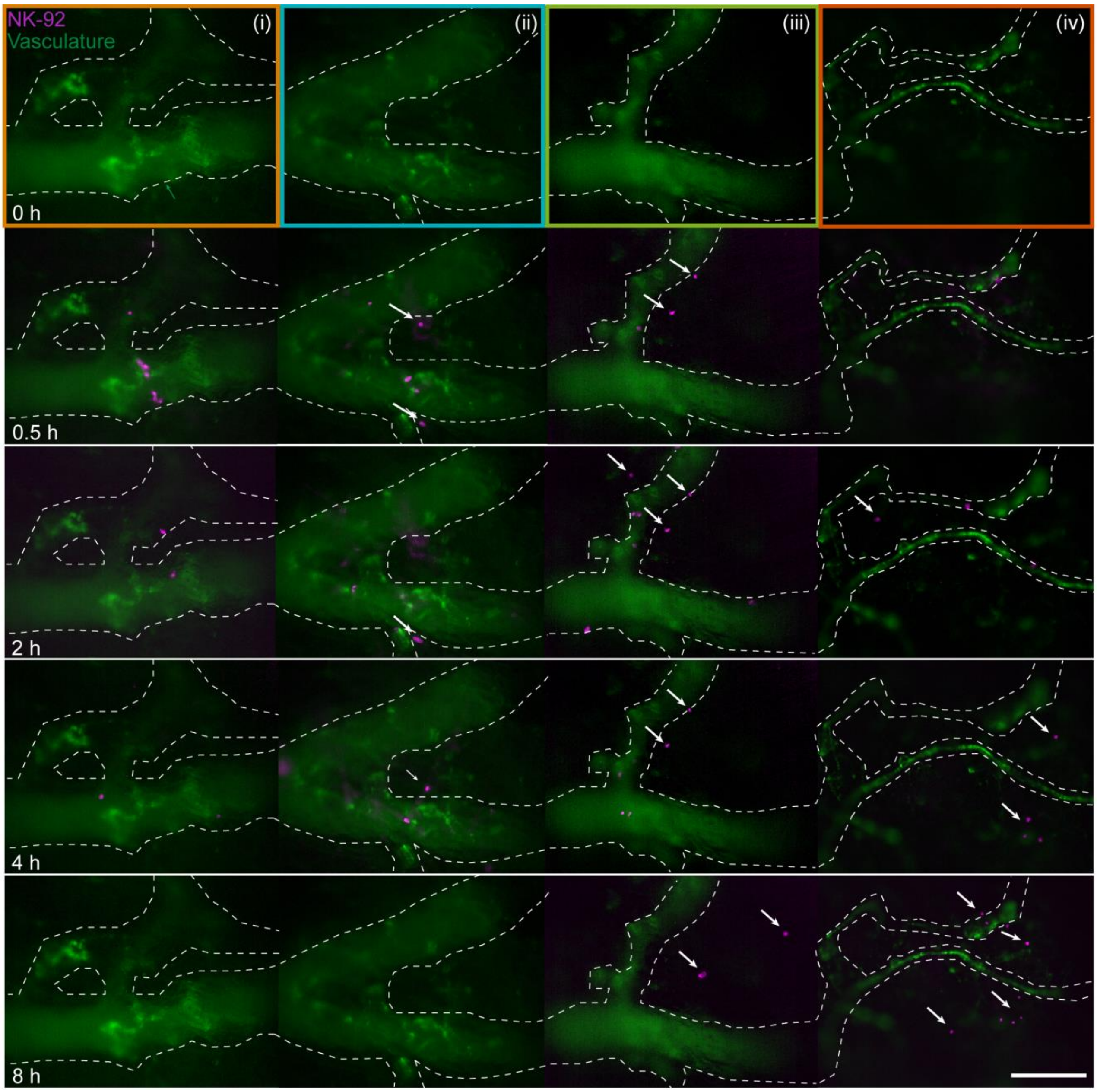

**Figure S29.** Representative dual-channel microscopic images recorded from the tumor vasculature at different times and in different regions before and after NK-92 cells transfusion. Regions (i): yellow rectangle; (ii): blue rectangle; (iii): green rectangle; (iv): orange rectangle. Vasculature: green; NK-92 cells: magenta. NK-92 cells infiltrating into the tumor parenchyma: white arrows. Scale bars, 50 μm.

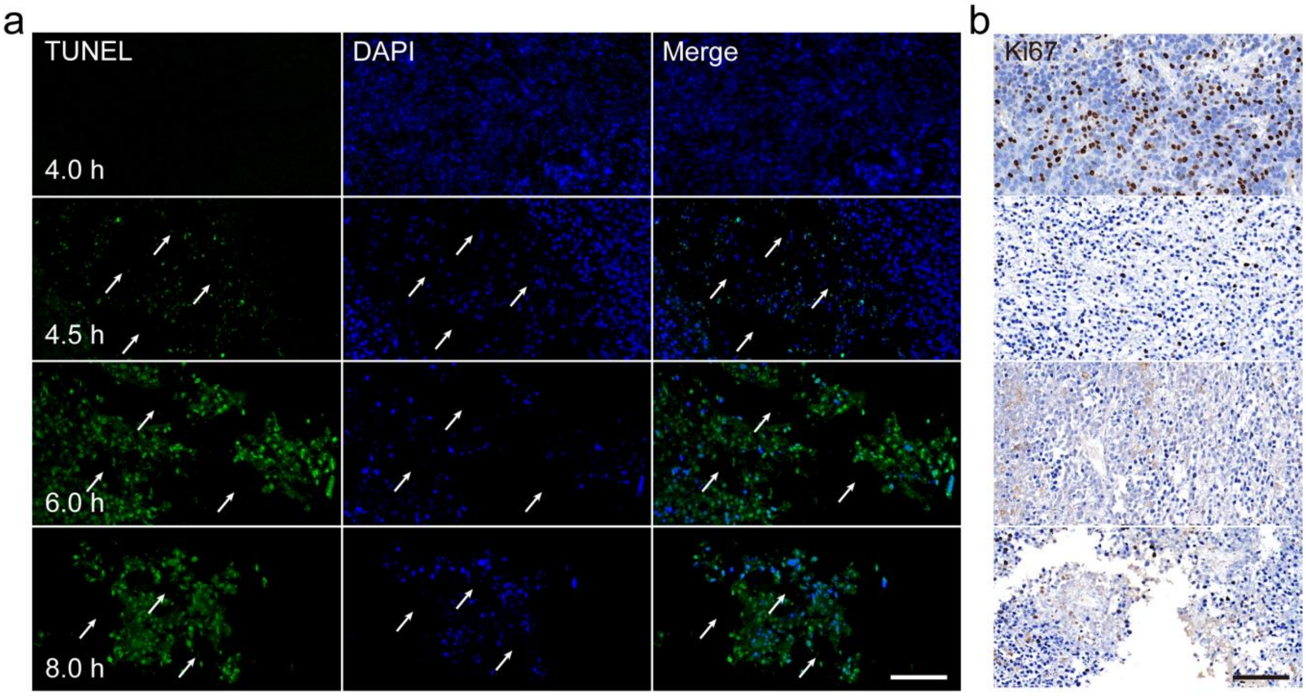

**Figure S30.** (**a**) TUNEL immunofluorescence was used to detect apoptosis in mouse tumor tissues induced by immunocytotherapy. White arrows represented detached tumor parenchyma. Scale bar, 100 μm. (**b**) Ki67 immunohistochemistry assessed the inhibitory effects of immunocytotherapy on tumor cell proliferation in mice. Scale bar, 100 μm.

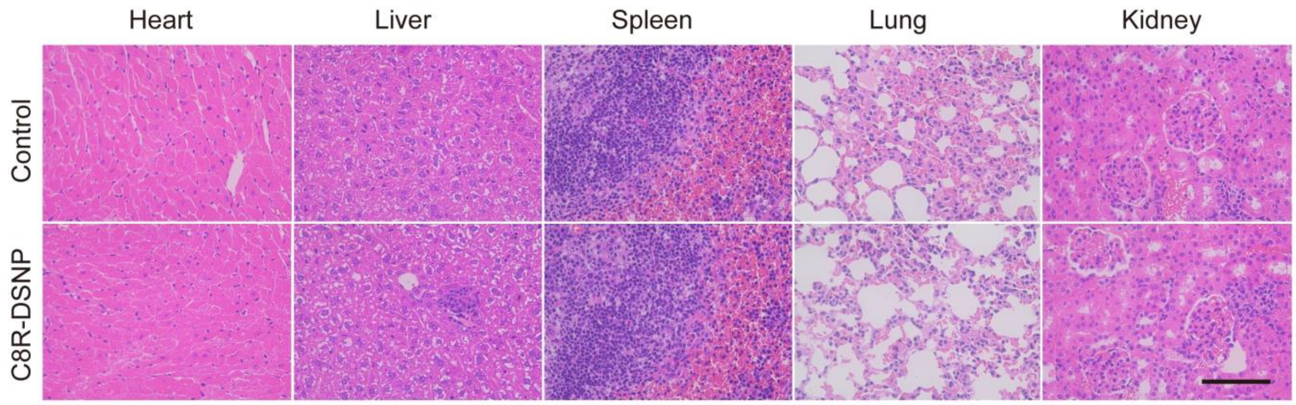

**Figure S31.** Histological studies of hematoxylin and eosin (H&E). H&E staining of internal organs (heart, liver, spleen, lungs, and kidneys) was obtained at 24 h post-injection of C8R-DSNP (200 μL, 500 μM) via tail vein. These results indicate that C8R-DSNP has low toxicity and does not induce pathological changed. Scale bar, 200 μm.

**Table S1.** Routine blood examination index in healthy mice and mice injected with C8R-DSNP

| Parametric | Results | Unit | Reference range |
|---|---|---|---|
| WBC | 1.2 | 10^9/L | 0.8-10.6 |
| Lymph# | 0.9 | 10^9/L | 0.6-8.9 |
| Mon# | 0.1 | 10^9/L | 0.04-1.4 |
| Gran# | 0.2 | 10^9/L | 0.23-3.6 |
| Lymph% | 75.2 | % | 40-92 |
| Mon% | 4.5 | % | 0.9-18 |
| Gran% | 20.3 | % | 6.5-50 |
| RBC | 7.64 | 10^12/L | 6.5-11.5 |
| HGB | 132 | g/L | 110-165 |
| HCT | 25.8 | % | 35-55 |
| MCV | 33.9 | fL | 41-55 |
| MCH | 17.2 | pg | 13-18 |
| MCHC | 511 | g/L | 300-360 |
| RDW | 32.0 | % | 12-19 |
| PLT | 429 | 10^9/L | 400-1600 |
| MPV | 5.6 | fL | 4.0-6.2 |
| PDW | 15.5 | | 12.0-17.5 |
| PCT | 0.283 | % | 0.100-0.780 |

#### Author Contributions

F.Z. developed the original concept. F.Z. and Y.F. designed and directed the study. L.H. and J.M. contributed equally to this study. L.H. and J.M. performed the nanosensor synthesis, characterization, and optical imaging experiments, and J.W. participated in the characterization experiments. L.H., T.Z., and B.Y. performed cellular and animal experiments. F.Z., Y.F., L.H., J.M. and A.L. analyzed results and wrote the manuscript. L.H. and J.M. analyzed the results and wrote the manuscript, graphs and supplementary information. All authors participated in the discussion and editing of the manuscript.
